# Actomyosin forces trigger a conformational change in desmoplakin within desmosomes

**DOI:** 10.1101/2024.11.19.624364

**Authors:** Yinchen Dong, Ahmed Elgerbi, Bin Xie, John S. Choy, Sanjeevi Sivasankar

## Abstract

Desmosomes are essential cell-cell adhesion organelles that enable tension-prone tissue, like the skin and heart, to withstand mechanical stress. Desmosomal anomalies are associated with numerous epidermal disorders and cardiomyopathies. Despite their critical role in maintaining tissue resilience, an understanding of how desmosomes sense and respond to mechanical stimuli is lacking. Here, we use a combination of super-resolution imaging, FRET-based tension sensors, atomistic computer simulations, and biochemical assays to demonstrate that actomyosin forces induce a conformational change in desmoplakin, a critical cytoplasmic desmosomal protein. We show that in human breast cancer MCF7 cells, actomyosin contractility reorients keratin intermediate filaments and directs force to desmoplakin along the keratin filament backbone. These forces induce a conformational change in the N-terminal plakin domain of desmoplakin, converting this domain from a folded (closed) to an extended (open) conformation. Our findings establish that desmoplakin is mechanosensitive and responds to changes in cellular load by undergoing a force-induced conformational change.

## Introduction

Desmosomes are critical cell-cell junctions that promote adhesion between cells and preserve tissue integrity. They enable tension-prone tissues like the skin and heart to withstand mechanical stress [1]. Desmosomal anomalies are associated with numerous diseases such as epidermal autoimmune disorders and arrhythmogenic ventricular cardiomyopathy [2–4]. Despite their critical role in maintaining tissue mechanical resilience, an understanding of how desmosomes respond to mechanical stimuli is lacking.

Desmosomes are composed of transmembrane desmosomal cadherins, desmoglein (Dsg) and desmocollin (Dsc), and cytoplasmic plaque proteins: plakophilin, plakoglobin, and desmoplakin (DP). The extracellular regions of the cadherins from opposing cells interact to mediate adhesion, while the plaque proteins bind to the cadherin cytoplasmic tail (Fig. 1A) [5]. DP anchors desmosomes to the keratin intermediate filament (KIF) cytoskeleton, and this linkage is essential for maintaining tissue integrity and facilitating intracellular signal transduction [6, 7]. Several studies show that KIFs work in cooperation with the actomyosin cytoskeleton to modulate the mechanical state of desmosomes [8–10]. Disassociating KIFs from circumferential actin belts alters tension across desmosomes [8]], while disassembling the actin cytoskeleton decreases desmosome resilience [9]. However, the molecular mechanisms by which actomyosin forces alter desmosome structure and function are not understood.

**Fig. 1:**
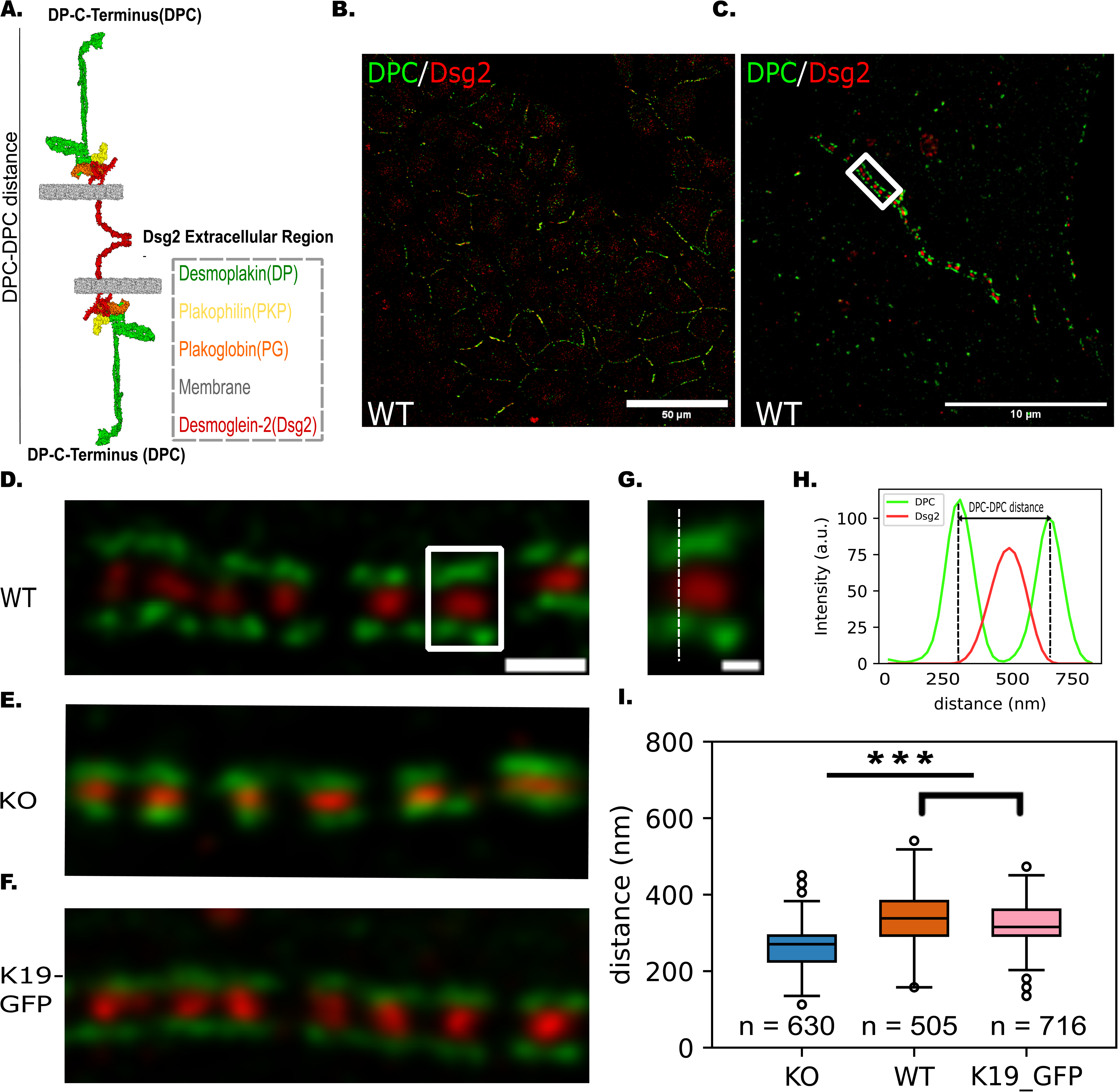
Keratin-19 influences desmosome architecture. **(A)** Schematic of desmosome structure. **(B)** Representative confocal image of desmosome puncta along cell-cell borders in the WT cells. The desmosomes were immunolabeled for DPC (green) and Dsg2 (red). Scale bar: 50 μm. **(C)** Representative STED image of desmosomes showing a characteristic ‘railroad track’ pattern. Scale bar: 10 μm. **(D-F)** Representative close-up STED images of desmosomes exhibiting the signature ‘railroad track’ pattern in **(D)** MCF7 WT cells, **(E)** MCF7 K19-KO cells, and **(F)** MCF7 K19-GFP cells. Scale bar: 500 nm. **(G)** The boxed area from the representative STED images of WT cells shows a closer look at an individual DP railroad track with Dsg2 between the two parallel DP plaques. Scale bar: 200 nm. **(H)** Line-scan analysis of DPC and Dsg2 fluorescence intensity (indicated by the dashed line in E). **(I)** Quantification of DPC-DPC distance from WT, K19-GFP, and K19-KO cells shows that desmosomes in WT and K19-GFP cells are wider than in K19-KO cells. In all boxplots, the box represents the 25th and 75th percentiles with the median indicated, and whiskers reach 1.5 times the interquartile range (IQR), defined as the difference between the 25th and 75th percentiles. Data points outside the whiskers are shown as outliers. Number of data points (n) = 630 (WT), 505 (K19-KO), 716 (K19-GFP) (n represents the number of line scans across the desmosomes); Number of replicates (N) = 3. Kruskal-Wallis Test, followed by Dunn’s multiple comparison Test; ***, P<0.001.

Disruption of the linkage between DP and KIF impairs intercellular adhesion and compromises tissue integrity [10, 11]. Knocking-out keratin leads to elevated phosphorylation of DP, resulting in accelerated endocytosis of desmosomes and rupture of epithelial sheets under rotational stress [12]. Weakening connections between DP and KIF networks dramatically alters the mechanical properties of cells, such as cell stiffness and intercellular forces [9]. While these data suggest that DP is crucial in regulating the response of desmosomes to mechanical stress, the biophysical mechanism by which DP mechanically regulates the desmosome is unknown.

The structure of DP offers hints into its mechanoregulatory function. DP is a large, 2871 amino acid protein composed of an N-terminal region, a central rod domain that facilitates DP homodimerization [3], and a C-terminal region that is crucial for aligning and binding KIFs [13]. The N-terminal region of DP is subdivided into a globular head domain and a plakin domain, each playing distinct roles [14]. The globular head domain binds to the C-termini of desmosomal cadherins and recruits plakophilin and plakoglobin [15]. The plakin domain, characterized by six spectrin repeat domains interspersed with Src homology domains, exhibits structural flexibility, as observed in a previous *in vitro*, cell-free small angle X-ray scattering study [16]. That investigation revealed that the plakin domain adopts either a folded “U” shape or a fully extended “I” shape, indicating a significant degree of structural adaptability [17]. This finding suggests that actomyosin forces could induce a conformational change in the N-terminal region of DP, thereby offering a molecular pathway for DP mechanosensation. However, force-induced conformational changes in DP, particularly within the cell, have never been demonstrated.

DP exists in two major splice isoforms, I and II, which differ only in the length of their rod domain. Our study focuses on DP isoform I (DPI) which is expressed in high levels in epithelia and cardiomyocytes [18]. We demonstrate, in human breast cancer MCF7 cells, that actomyosin forces induce a conformational change in the N-terminal region of DPI. Since MCF7 cells are highly enriched in keratin 19 (K19), we interrogated wild type (WT), K19-knock out (K19-KO), and K19-GFP rescue (K19-GFP) cells with super-resolution microscopy, FRET-based tension sensors, pharmacological inhibition and activation of actomyosin contractility, co-immunoprecipitation (co-IP), and atomistic Molecular Dynamics (MD) simulations. We demonstrate that actomyosin contractility reorients the KIF and directs force to DPI along K19 filaments. We show that these forces induce a conformational change in the N-terminal plakin domain, from a folded ‘closed’ to an extended ‘open’ conformation. Our findings establish, at the molecular level, that DP is a mechanosensor within the desmosome.

## Results

### K19 remodels desmosome architecture

Desmosomes are characterized by distinct “railroad” track structures, represented by discrete parallel fragments of DP, which sandwich the desmosomal cadherins [19]. To validate this signature structure of desmosomes in MCF7 cells, we conducted immunostaining targeting the extracellular region of isoform 2 of desmoglein (Dsg2) and the C-terminal region of DPI (DPC) using confocal and super-resolution Stimulated Emission Depletion (STED) microscopy. Confocal imaging revealed co-localization of DPC and DSG2 signals, indicating the formation of desmosome puncta along cell–cell junctions (Fig. 1B). STED imaging in WT cells showed well-defined DP “railroad” tracks, as depicted in Figs. 1C-D. To investigate the effect of K19 on desmosome architecture, we used the K19-KO cell line [20]. Successful ablation of K19 in these cells was confirmed using western blotting (Fig. S1). Surprisingly, in the K19-KO cells, STED imaging revealed more compact desmosomes (Fig. 1E). To quantify the width of the desmosomes, we measured the distance between opposing DP plaques in the STEAD images by drawing a line scan across the desmosome complex (Fig. 1G) and measuring the distance between the two peaks of DPC signal (designated as DPC-DPC distance), as illustrated in the fluorescence intensity plot (Fig. 1H). This quantitative analysis confirmed that desmosomes in K19-KO cells were significantly narrower than desmosomes in WT MCF7 cells (334 ± 61 nm for WT cells and 264 ± 65 nm for K19-KO cells; Fig. 1I).

To corroborate that the observed shift in desmosome width was directly related to the absence of K19, we stably rescued the K19-KO cells with EGFP-tagged K19 (K19-GFP cells). Western blots validated the successful expression of K19-GFP in the rescued cells (Fig. S2). STED imaging showed that desmosome width in the K19-GFP cells recovered to a level similar to the WT cell line (322 ± 56; Figs. 1F & 1I). Together, these findings demonstrate that desmosome architecture in MCF7 cells is strongly correlated to the presence/absence of K19. In the presence of K19, the desmosome width is larger, while the desmosome narrows when K19 is knocked out.

### The interplay between K19 and the actomyosin networks governs desmosome width

The cytoplasmic KIF network binds to DP at its C-terminus (illustrated in Fig. 2A) and interactions between the KIF network and elements of the actomyosin cytoskeleton have been observed at cell-cell junctions and the cell-substrate interface [8, 21]. To investigate how changes in the KIF network contribute to desmosome remodeling, we first inspected the intracellular localization of K19, DP, and F-actin filaments in K19-GFP cells. K19 was found to be colocalized with the DP “railroad” tracks along the cell border (Fig. 2B). Additionally, actin filaments were observed to align with the K19 filaments (Fig. 2B). Based on the colocalization of K19 with DP and the coalignment of K19 with F-actin filaments, we hypothesized that K19 interacts with both DP and F-actin filaments.

**Fig. 2:**
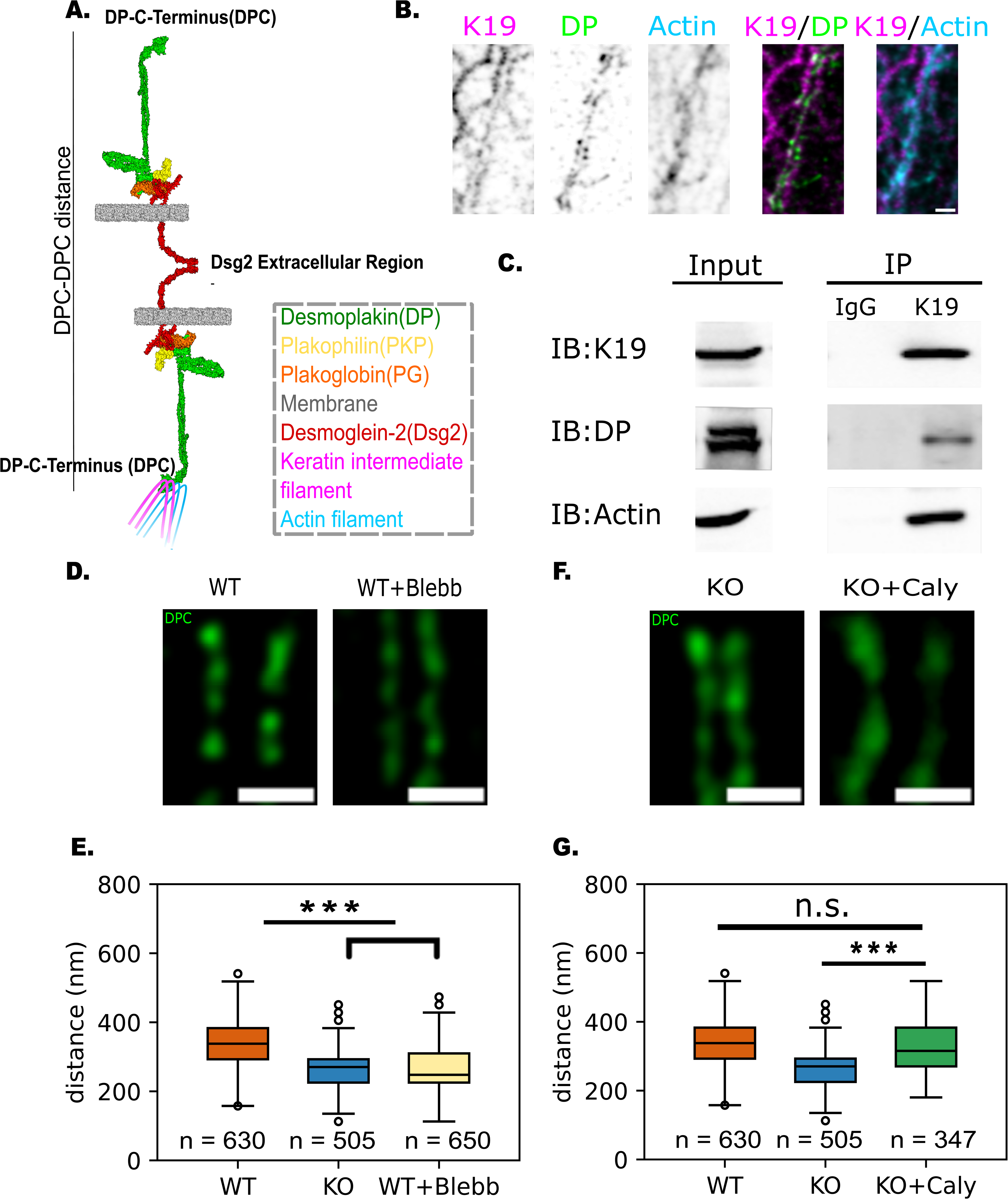
Interplay between keratin-19 and actomyosin network determines desmosome width. **(A)** Schematic of the interactions between the desmosome, keratin intermediate filaments, and actin filaments. Keratin filaments bind to the C-terminal of DP and interacts with actin filaments in the cytoplasm. **(B)** Representative STED images of K19-GFP rescue cells immunolabeled for K19 (magenta), DP (green), and actin (cyan). The individual components are shown in grayscale to highlight the structural features. The merged images between K19 and DP or actin are shown at the right. Scale bar: 500 nm. The images show that K19 filaments colocalize with the DP “railroad” tracks and with cortical actin belts. **(C)** Co-IP in WT cells performed with anti-K19 antibody or IgG control. The Co-Ips show that K19 interacts with both DP and actin. **(D)** Representative STED images of DPC (green) at the cell border of WT cells and Blebbstatin-treated WT cells (WT+Blebb). **(E)** Quantification of desmosome widths from WT, K19-KO, and WT+Blebb cells showing that Blebbistatin treatment reduces the desmosome width to a similar level as in the K19-KO cell line. n = 630 (WT), 505 (K19-KO), 650 (WT+Blebb); N = 3. Kruskal-Wallis Test, followed by Dunn’s multiple comparison Test; ***, P<0.001. **(F)** Representative STED images of DPC (green) at the cell border of K19-KO cells and Calyculin A-treated K19-KO cells (KO+Caly). **(G)** Quantification of desmosome widths from WT, K19-KO, and KO+Caly cells showing that Calyculin A increases the desmosome width of K19-KO cells to a similar level as in the WT cell line. n = 630 (WT), 505 (K19-KO), 347 (KO+Caly); N = 3. Kruskal-Wallis Test, followed by Dunn’s multiple comparison Test; ***, P<0.001.

To validate this hypothesis, we conducted Co-IP experiments using IgG as the control and K19 as the bait protein. The results revealed that K19 interacts with both DP and actin, consistent with our previous immunofluorescence microscopy observations (Fig. 2C). To confirm that the proteins pulled down with K19 were due to specific interactions, we performed a K19 pulldown in K19-KO cells. Neither DP nor actin filaments were detected in the pulldown from K19-KO cells (Fig. S3), supporting that their association with K19 in WT cells reflect specific interactions. To further confirm the specificity of K19 interactions with DP and actin, we also blotted for the membrane-associated protein E-cadherin in both WT and K19-KO cells. The absence of E-cadherin in the pull-down indicated that associations with DP and actin were not due to general membrane localization but are specific to K19 (Fig. S4). Finally, the Co-IP experiments showed that in addition to DP, the desmosomal plaque protein plakophilin-3 also interacted with K19 (Fig. S5), which further supports the interaction between K19 filaments and the desmosome complex.

It has previously been demonstrated that actin stress fiber formation strongly correlates with the presence of keratin filaments in cells [22]. We therefore used confocal fluorescence microscopy to image if actin stress fibers, which are a proxy for intracellular forces, are present in WT, K19-GFP, and K19-KO cells (Fig. S6A). Quantitative data analysis showed that compared to WT and K19-GFP cells, actin stress fibers in the K19-KO cells were dramatically decreased (Fig. S6B). This suggests that mechanical tension in the K19-KO cells is lower compared to WT and K19-GFP cells. We, therefore, hypothesized that actomyosin-generated forces could be transmitted to the desmosome complex through associations with the KIF network.

To test this hypothesis, we first used Blebbistatin, a myosin-II-specific ATPase inhibitor, to reduce the actomyosin contractility in the WT cell line (WT+Blebb) and measured the desmosome width. STED imaging showed more compact desmosomes in the WT+Blebb cells (Fig. 2D); desmosome width in WT+Blebb cells was reduced to a similar level as K19-KO cells (263 ± 62 for WT+Blebb; Fig. 2E). At the same time, the control group treated with dimethyl sulfoxide (DMSO, Blebbistatin’s solvent), retained similar desmosome widths as the WT cells (Fig. S7).

Next, to test if stimulating actomyosin contractility increases desmosome width, we treated the K19-KO cells with Calyculin A, a phosphatase inhibitor (KO+Caly). STED imaging showed that the width of desmosomes in the KO+Caly cells increased to a level comparable to WT cells (Fig.2F). Quantitative analysis of the images showed that upon Calyculin A treatment, the desmosome width in K19-KO cells was 333 ± 83 (Fig. 2G). Taken together, these data demonstrate that the K19-dependent changes in desmosome architecture are a consequence of a mechanical pathway that is established due to interactions between K19, F-actin, and DP. Consequently, contractile actomyosin forces are transmitted to and reorganize the desmosome, resulting in an increase in desmosome width.

### The N-terminal plakin domain of DP transitions from a closed to open conformation when pulled

Since previous studies have reported structural flexibility in the N-terminal plakin domain of DP [17], we hypothesized that the increase in desmosome width in WT cells was due to a force-induced conformational change in this region of DP (Fig. 3A).To test this hypothesis, we specifically stained both the DPI N-terminal region (DPN) on its globular head domain (which is upstream of the plakin domain) and the extracellular region of Dsg2. Using super-resolution imaging, wider desmosomes were observed in WT cells stained with DPC and DSG2 compared to K19-KO cells (Fig.3B). Interestingly, when stained with DPN and DSG2, both cell lines exhibited similar desmosome widths (Fig.3C). To gain a clearer understanding of the DP structural changes in WT and K19-KO cell lines, we measured the distances between Dsg2 and DPC, as well as Dsg2 and DPN within the half-unit of the desmosome complex. This analysis showed that while Dsg2-DPC distance was significantly greater in WT compared to the K19-KO cells (167 ± 47 for WT cells and 132 ± 50 for K19-KO cells; Fig. 3D), Dsg2-DPN distance was similar in both cell lines (101 ± 48 for WT cells and 99 ± 53 for K19-KO cells; Fig. 3E). As seen in Fig. 3G, the DP length (defined as the distance between DPN and DPC) was 33 nm greater in the WT cells compared to the K19-KO cells, suggesting that DP was elongated by ∼33 nm in the WT cell lines due to higher mechanical tension in these cells.

**Fig. 3:**
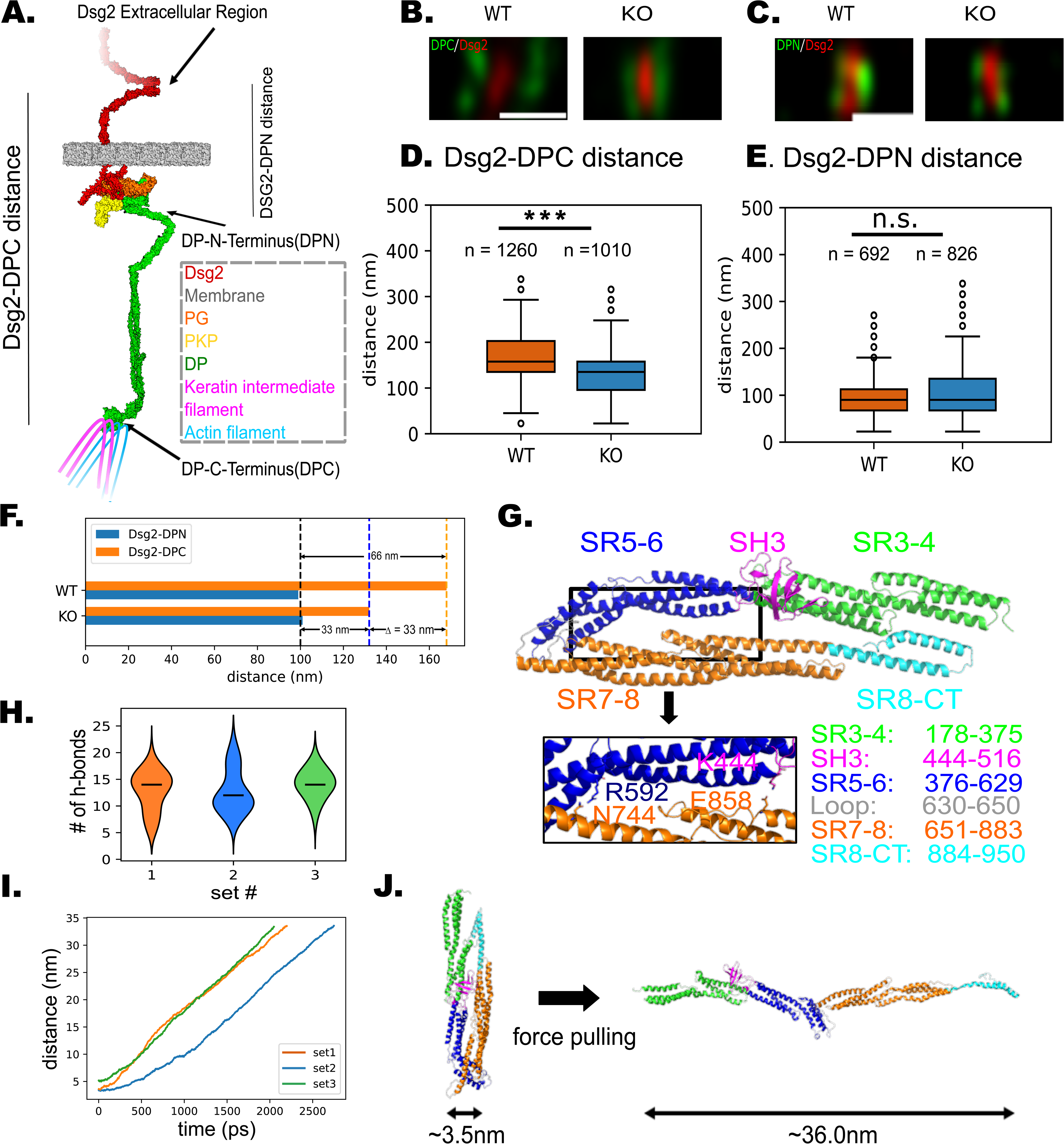
Desmoplakin undergoes force-induced conformational change in the plakin domain. **(A)** Schematic of the desmosome under tension. The N-terminal of DP transitions from a closed conformation to an open conformation. **(B)** Representative STED images of DPC (green) and DSG2 (red) at the cell border for WT and K19-KO cells. Scale bar: 500 nm. **(C)** Representative STED images of DPN (green) and DSG2 (red) at the cell border for WT and K19-KO cells. Scale bar: 500 nm. **(D)** Quantification of desmosome half-unit widths (Dsg2-DPC distance) from WT and K19-KO cells. n = 1260 (WT), 1010 (K19-KO); N = 3. Mann-Whitney’s U test; ***, P<0.001. The distances between Dsg2 and DPC are significantly greater in the WT compared to the K19-KO cells. **(E)** Quantification of desmosome half-unit widths (Dsg2-DPN distance) from WT and K19-KO cells. n = 692 (WT), 826 (K19-KO); N = 3. Mann-Whitney’s U test; ns, P>0.05. The Dsg2-DPN distance in both cell lines are similar. **(F)** The mean values from the results in D and E are summarized in the bar chart to compare the DP length (DPN-DPC distance) between the WT and K19-KO cells. The DP length is 66 nm for the WT and 33 nm for the K19-KO, indicating that DP extends 33 nm in the WT cells. **(G)** The entire plakin domain structure predicted by Alphafold2 suggests a U-shape conformation in the absence of force. Two common electrostatic interactions (R592-N744 and K444-E858) observed in all 3 MD simulations (below) are shown. **(H)** During the MD simulations, 10 to 15 hydrogen bonds were formed between the plakin domain’s long arm (SR3-4 and SR5-6) and short arm (SR7-8 and SR8-CT). **(I)** During the SMD simulations, the distance between the plakin domain’s C-terminus and N-terminus increased by 30∼33 nm. This elongation correlates with the distance change experimentally measured in Fig 3F. **(J)** Comparison of the plakin domain structure at the start (left) and at the end (right) of the SMD simulation. The structures show that the plakin domain was elongated by the pulling force without significant secondary structure unfolding.

To resolve the mechanistic basis by which DP transitions between open and closed structures, we proceeded to atomistically simulate the effect of force on the conformation of the highly conserved plakin domain of DP (residues 178-950). We first generated the entire plakin domain structure using Alphfold2 [23]. The predicted structure exhibited a ‘U-shaped’ folded conformation, with spectrin repeat (SR) domains SR3-6 and Src homology (SH) domain SH3 forming a long arm that interacts with a short arm composed of SR7-8 and SR8-CT domains (Fig. 3F). This prediction agrees well with the crystal structure of the long arm of the plakin domain [24] (RMSD ∼1.7Å, Fig. S8).

To identify the interactions crucial for stabilizing the folded conformation, we performed three force-independent molecular dynamics (MD) simulations on the predicted structure. In the MD simulations, the plakin domain stabilized within 20 ns (Fig. S9). During the simulations, 10-15 hydrogen bonds were observed between the long arm and the short arm, and the interactions mainly were observed between SR5-6 and SR7-8 regions (Fig. 3H and Fig. S10). Notably, two consistent salt bridge/hydrogen bond interactions, between four conserved amino acids, were observed across all three simulations: R592-N744 and K444-E858 (Fig. 3F).

To test if the folded plakin domain structure can be unfolded using pulling forces, we performed steered molecular dynamics (SMD) simulations (Video S1). In the SMD simulations, we fixed the position of the N-terminal residues and pulled the C-terminal regions (residues 919-950) using a constant force. Our simulations showed that the plakin domain’s long and short arms separated due to the pulling force without any secondary structure unfolding (Fig. 3J). Consequently, the plakin domain transitioned from the initial closed (Fig. 3J, left) to the final open (Fig. 3J, right) conformation (Video S1). In the open conformation, the distance between the plakin domain’s C-terminus and N-terminus increased by 33 nm compared to the closed structure (Fig. 3I), which is consistent with the change in DPC-DPC distance observed in the STED images (Fig. 3G).

### Keratin intermediate filaments reorient in response to changes in actomyosin forces

Given that actomyosin forces are transmitted to desmosomes via KIFs, a key question is how this force transmission is affected by the organization of keratin filaments. A previous study in *Xenopus laevis* showed that the KIF network reorganizes in response to force transmission across cell-cell contacts and becomes aligned with the direction of protrusive activity and tissue movement [25]. This raises the question as to whether a similar phenomenon occurs in MCF7 cells (Fig. 4A).

**Fig. 4:**
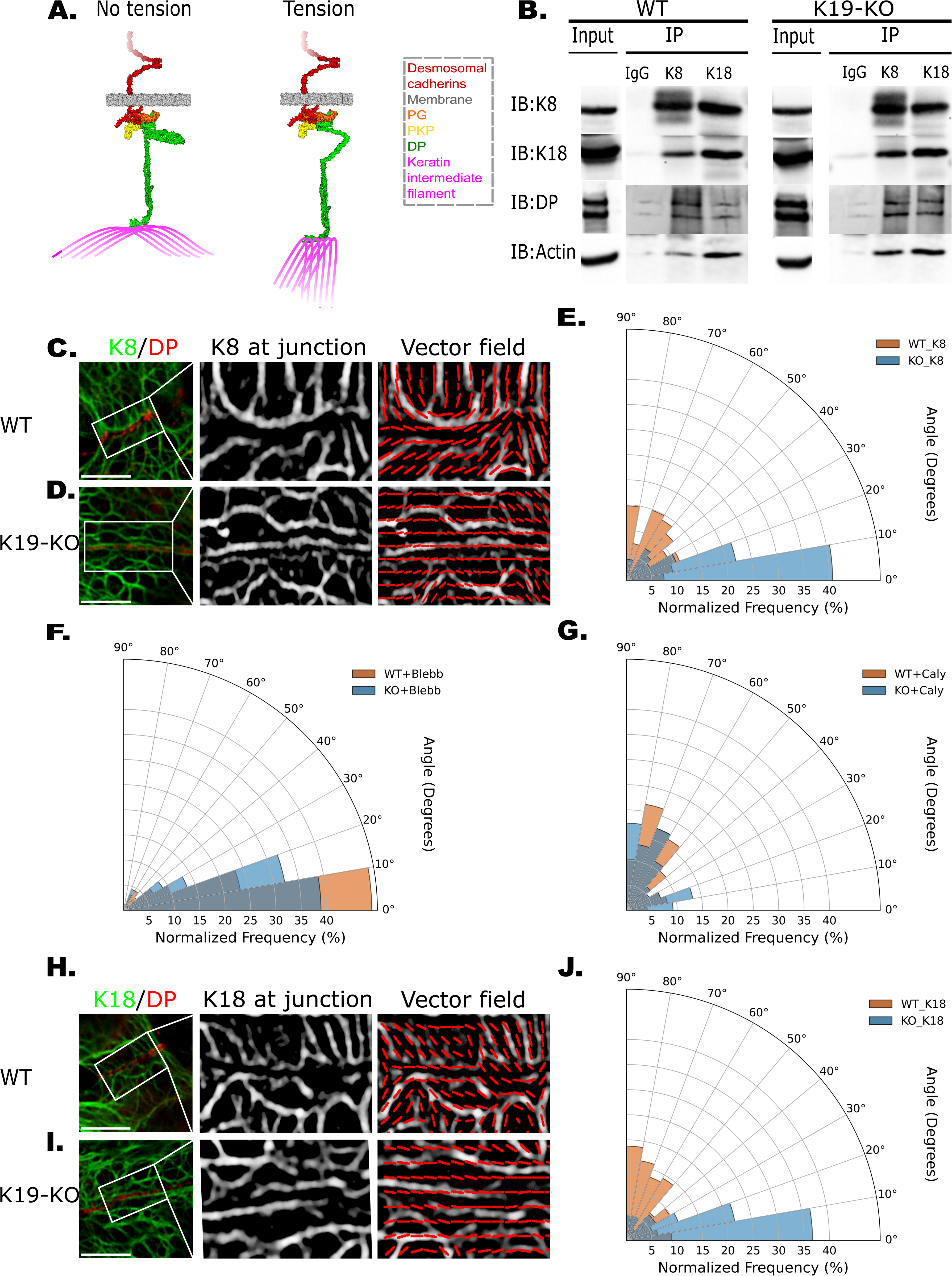
Keratin filaments reorient in response to changes in actomyosin forces: **(A)** Schematic representation of DP-associated keratin filaments at the cell-cell junction. Under no-tension conditions, DP-associated filaments are primarily aligned horizontally. Under tension, they transition to a radially organized structure that exerts force on the desmosome. **(B)** Co-IP of K8 and K18 in WT and K19-KO cells. Co-IP was performed using anti-K8, anti-K18 antibodies, or an IgG control. The Co-IPs reveal that both K8 and K18 interact with DP and actin. **(C-D)** Representative confocal images of WT **(C)** and K19-KO **(D)** cells immunolabeled for K8 (green) and DP (red). Scale bar: 5 µm (left panel). A magnified grayscale image highlights DP-associated K8 filaments at the junction with enhanced contrast (middle panel). The local dominant orientation of K8 filaments represented as a vector field image (right panel). **(E)** Polar histogram of the global dominant orientation of K8 filaments in WT and K19-KO cells. n = 108 (WT), 96 (K19-KO); N = 3. Mann-Whitney U test, P < 0.0001. The K8 filaments in WT cells exhibit a more radial organization, while K19-KO filaments are horizontally aligned. **(F)** Polar histogram of K8 filament orientation in WT and K19-KO cells treated with Blebbistatin. n = 90 (WT), 90 (K19-KO); N = 3. Mann-Whitney U test, P > 0.05. Following Blebbistatin treatment, WT filaments shift from a radial to a horizontal organization, making their global dominant orientation indistinguishable from K19-KO cells. **(G)** Polar histogram of K8 filament orientation in WT and K19-KO cells treated with Calyculin A. n = 98 (WT), 98 (K19-KO); N = 3. Mann-Whitney U test, P > 0.05. After Calyculin A treatment, K19-KO K8 filaments shift from a horizontal to a radial organization, making their global dominant orientation indistinguishable from WT cells. **(H-I)** Representative confocal images of **(H)** WT and **(I)** K19-KO cells immunolabeled for K18 (green) and DP (red). Scale bar: 5 µm (left panel). Magnified images highlighting K18 filaments at the junction and local dominant orientation of K18 filaments are shown in the middle and right panel. **(J)** Polar histogram of the global dominant orientation of K18 filaments in WT and K19-KO cells. n = 112 (WT), 120 (K19-KO); N = 3. Mann-Whitney U test, P < 0.0001. The global dominant orientation of K18 filaments in WT cells is significantly different from that in K19-KO cells, with WT filaments adopting a radial organization while K19-KO filaments remain horizontal, mirroring the pattern observed for K8 filaments.

In epithelial cells, KIFs are composed of heterodimers between type I and type II keratins, which assemble into antiparallel tetramers [26]. A previous RNA-sequencing study revealed that type II Keratin-8 (K8) was the most highly expressed isoform in the MCF7 cells, followed by roughly equal amounts of type I Keratin-19 (K19) and type I Keratin-18 (K18) [27]. This suggests that the primary keratin pairs in MCF7 cells are K8/K19 and K8/K18. Notably, K8 and K18 expression levels remained unchanged when K19 is knocked out (Fig. S1), which allowed us to stain K8 and K18 to investigate KIF organization. When we performed Co-IP experiments using K8 and K18 as bait, we observed that both keratin isoforms interacted with DP and actin filaments in WT and K19-KO cells (Fig. 4B). This suggested that K8/K18 filaments could also transmit actomyosin forces to desmosomes.

We first imaged the organization of K8 filaments in WT and K19-KO cells by staining both K8 and DP. In WT cells, the K8 filaments at junctions exhibited a more radial organization (Fig. 4C), whereas, in K19-KO cells, the K8 filaments were primarily arranged horizontally (Fig. 4D). We quantified filament organization by determining their local orientation, defined as the angle of the most dominant filament direction with the primary axis of the cell junction (which was set as 0°), and aggregated the local orientations into a polar histogram (Fig. 4C-D). Our quantitative analysis showed that over 60% of WT K8 filaments had a dominant orientation between 40° and 90°, whereas more than 60% of K19-KO K8 filaments were oriented between 0° and 20° (Fig. 4E).

When actomyosin force was inhibited via Blebbistatin treatment, the K8 filament orientation in the WT cells changed dramatically with over 70% of filaments oriented between 0° and 20° (Fig. 4F). In contrast, K19-KO cells treated with Blebbistatin showed no significant change in K8 filament orientation (Fig. 4F). On the other hand, when actomyosin contractility in the cells was enhanced using Calyculin A treatment, K8 filament organization in K19-KO cells changed dramatically with over 60% of filaments having a dominant filament orientation between 40° and 90° (Fig. 4G). In contrast, K8 filament orientation in WT cells treated with Calyculin A remained largely unchanged (Fig. 4G). At the same time, the control group treated with Blebbistatin and Calyculin A’s solvent, DMSO, retained similar keratin organizations as the WT and K19-KO cells (Fig. S11).

Next, to distinguish between K8/K19 and K8/K18 filaments, we also imaged K18 filament organization in WT and K19-KO cells. As observed in our K8 filament images, staining for K18 and DP revealed more radially organized filaments in WT cells and horizontally organized filaments in K19-KO cells (Fig. 4H-J). Taken together, these findings indicated that KIFs reorganize in response to changes in actomyosin forces, shifting from predominantly ‘poor force-transmitting’ horizontal orientation in a low-tension condition to a more ‘efficient force-transmitting’ radial organization under load. Consequently, K19-KO and WT+Blebb cells exhibit horizontally oriented KIF network and more compact desmosomes while WT cells and KO+Caly display a radially organized KIF network and wider desmosomes.

### Desmoplakin is mechanically loaded in WT cells

Next, we proceeded to directly compare tension across DP in WT and K19-KO desmosomes using a recently published DPI Förster Resonance Energy Transfer (FRET) tension sensor [28]. This F40-based tension sensor (TS), is comprised of a tension-sensing linker peptide (GPGGA)₈ flanked by the donor fluorophore mTFP1 and the acceptor fluorophore mEYFP, that is inserted into an unstructured region following the rod domain, just before Pro1946 in DPI (Fig. 5A) [28]. Additionally, we used a no-tension control consisting of a truncated DP that lacked the C-terminal keratin-binding domain to account for local environmental changes that may affect FRET efficiencies (Fig. 5A). While these truncated DP controls still localized to intercellular junctions, they were unable to interact with keratin filaments and thus could not bear forces [11, 28, 29]. Lastly, a donor-only control was also included to determine the FRET efficiency. We transiently expressed the DP constructs in WT and K19-KO cells and analyzed the fixed samples. DP-TS localized to cell-cell contacts, with distinct DP puncta observed in both WT and K19-KO cells (Fig. 5B). Similarly, the truncated DP control and donor-only control exhibited the expected subcellular localization at cell junctions, forming clear desmosomal puncta (Fig. S12). These results indicate that all three DP constructs were correctly localized to cell-cell contacts and successfully formed desmosomes, enabling the measurement of DP tension at the desmosome.

**Fig. 5:**
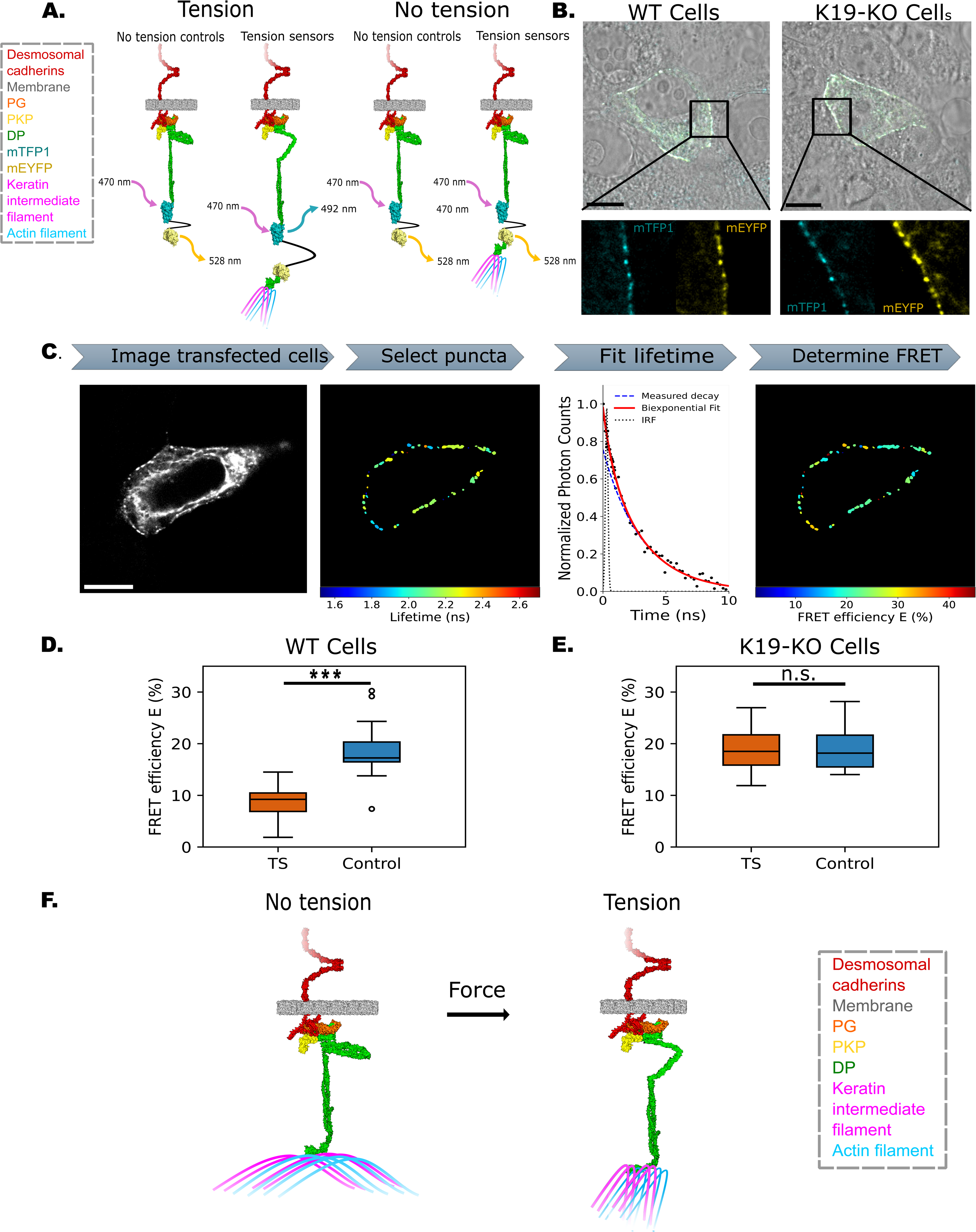
Desmoplakin is mechanically loaded in WT cells but not in K19-KO cells. **(A)** Schematic illustration of the DP FRET tension sensor (DP-TS) and no-tension control. The tension-sensing module consists of a linker peptide flanked by mTFP1 (donor fluorophore) and mEYFP (acceptor fluorophore). In the absence of tension, the FRET pair stays together in both constructs. Under tension, the FRET pair is pulled apart in DP-TS, but remains together in the no-tension control which lacks the keratin-binding C-terminal region. **(B)** Representative STED images of WT and K19-KO cells transfected with DP-TS. The overlay of fluorescent signals with the bright-field image shows the proper localization of DP-TS at the cell-cell contacts in both WT and K19-KO cells. Scale bar: 10 µm. The zoomed-in images reveal the formation of distinct DP puncta along the cell border. **(C)** Illustration of the workflow for FRET-FLIM analysis. First, lifetime data images were acquired from transfected cells in fixed samples. Scale bar: 10 µm. Next, individual DP puncta were manually selected as regions of interest (ROIs) along the cell border. The fluorescence lifetime for each DP puncta was determined by fitting a bi-exponential model to distinguish true lifetime measurements from the effects of the instrument response function (IRF). Finally, FRET efficiency (E) was calculated from the lifetime measurements as described in the methods. **(D)** Quantification of FRET efficiency for DP-TS and no-tension control constructs in WT cells. The median FRET efficiency of the tension sensor was significantly lower than the control, indicating that DP is mechanically loaded in WT cells. *n* = 30 images, *N* = 3, *P* < 0.0001. **(E)** Quantification of FRET efficiency for DP-TS and no-tension control constructs in K19-KO cells. The median FRET efficiency of DP-TS was indistinguishable from that of the control, suggesting that DP experiences undetectable tension in K19-KO cells. *n* = 30 images, *N* = 3, *P* > 0.05. **(F)** Model for force-induced DP conformational change in desmosomes. When desmosomes experience no tension, DP-associated keratin intermediate filaments align horizontally with the cell membrane, maintaining a closed DP conformation. Under mechanical stress, the keratin filaments rearrange into a more radial orientation, effectively transmitting forces to DP. This results in a conformational change in the flexible DP plakin domain, converting this domain from a folded (closed) to an extended (open) conformation.

Next, we quantified DP-TS FRET efficiencies using fluorescence lifetime imaging microscopy (FLIM) in confluent WT and K19-KO monolayers following the workflow illustrated in Fig. 5C. When imaging each transfected cell, we first isolated individual DP puncta and measured their lifetime using a bi-exponential fit, accounting for the effects of the instrument response function (IRF) and autofluorescence, which yielded short lifetime measurements. Finally, FRET efficiency was determined based on these lifetime measurements. In the WT cells, the FRET efficiency was significantly lower in the TS (8.80% ± 2.76%) compared to the control (18.5% ± 4.46%), indicating that DP in these cells experienced substantial mechanical loading (Fig.5D). In contrast, the FRET efficiency of TS (18.3% ± 3.96%) in K19-KO cells was similar to the control (19.0% ± 3.88%) (Fig. 5E), suggesting that no detectable tension was present across DP in the K19-KO cells. Based on the 1–6 pN sensitivity of the F40 tension sensor and its averaged FRET–force calibration curve [28, 30], we estimated that the tension in WT cells was close to the upper range, around 5 pN. These direct measurements of forces experienced by DP revealed high tension in WT cells and no detectable tension in K19-KO cells, consistent with our previous analysis of actin stress fibers.

## Discussion

Based on our data, we propose a model (Fig. 5F), where in the absence of tension, DP-associated keratin filaments align horizontally, maintaining a closed DP conformation. Under stress, these filaments adopt a radial orientation, transmitting forces to the desmosome and inducing a conformational shift in DP via its flexible plakin domain (Fig. 5F). Several recent studies have highlighted the potential mechanosensitive role of DP in desmosomes. These studies demonstrate that that DP experiences tension when cells are subjected to mechanical pulling [28] and show that the DP-KIF linkage is essential for resisting mechanical strain [9, 31]. Building on these findings, we demonstrate that DP functions as a mechanosensor by undergoing a conformational change in response to alterations in cytoskeletal tension. The DP conformational change described in our study is analogous to conformational changes in α-catenin, associated with classical cadherins in Adherens Junctions [32–34]. In the case of α-catenin, tension from the actin cytoskeleton induces a conformational change that opens its central M-domain and exposes cryptic binding sites for vinculin and other actin-binding proteins [32–34]. It is likely that the force-induced opening of DP also exposes cryptic binding sites for junctional proteins that signal and remodel the desmosome and intermediate filaments. The force-induced unfolding of DP could also serve as a damper of mechanical stress that maintains mechanical homeostasis within tissue.

Our data suggests that besides actomyosin forces, conformational changes in DP are also mediated by changes in keratin reorganization. Since Co-IP experiments suggest that both K8 and K18 associate with actin and DP, it is likely that K8/K18 filaments also transmit actomyosin forces to DP in the K19-KO cells. Why then do the DPs in the K19-KO cells remain in a closed conformation? Our data suggests that the loss of K19 alters the organization of keratin filaments, which in turn affects the efficiency of actomyosin force transmission to desmosomes. We show that in the K19-KO cells, K8/K18 filaments are oriented cortically, almost parallel to the desmosome, which result in tangentially actomyosin forces which cannot open DP. In contrast, since the KIFs are oriented radially in WT cells, desmosomes are exposed to normal forces which can induce a conformational change in DP. Indeed, using FRET tension sensors we demonstrate that DP in the WT cells are exposed to higher tension than DP in the K19-KO cells. We confirmed that the force-induced widening of desmosomes was not due to differences in the expression levels of desmosomal or cytoskeletal proteins between the two cell lines by comparing the amounts of essential desmosomal and cytoskeletal components in both WT and K19-KO cells. Immunoblotting results showed no significant differences in the levels of desmosomal cadherins (Dsg2, Dsc1, and Dsc2-3), plaque proteins (plakophilin-1, plakophilin-3, and DP), or cytoskeletal proteins, including K8 and K18, F-actin, and myosin heavy chain 10. These results indicate that the observed changes in desmosome organization under force are not due to altered protein expression levels but rather the result of modified mechanical conditions driven by the cooperative interactions between KIF and the actomyosin network.

Earlier studies using FRET-based tension sensors in resting MDCK cells revealed a small amount of tension across Dsg2 but not across DP [28, 35]. However, the Dsg2 FRET-based tension sensor showed a higher tension level in the contracting cardiac myocytes, suggesting that junctional tension depends on both cell type and the specific cellular environment [35]. To validate our analysis, we also measured DPC-DPC distances in resting MDCK cells. Our measurements revealed that the desmosome width in MDCK cells is comparable to that observed in K19-KO cells, implying that DP in resting MDCK cells indeed does not undergo conformational changes (Fig. S13). Additionally, FRET-based tension measurements in MCF7 cells, using the same DPI constructs as in previous studies, revealed high levels of tension across DP in WT cells but no detectable tension in K19-KO cells. These FRET-based tension measurements, along with the DPC-DPC distance data, support a correlation between desmosomal tension level and desmosome width, demonstrating that wider desmosomes result from higher tension across the desmosomes.

Previous rotary shadow imaging of DPI reported an average C-to-N-terminal distance of 162 nm [36]. In contrast, transmission EM imaging of desmosomes in bovine nasal epidermis reported a DP length of ∼42 nm [37] while super-resolution dSTORM microscopy in human epidermal keratinocytes measured DP lengths of ∼54 nm [19] which suggests that DP is tilted at an angle rather than being strictly perpendicular to the plasma membrane. In agreement, our STED measurements reveal a DPI length, based on the antibody-labeling, of approximately 66 nm in WT cells and 33 nm in K19-KO cells. Importantly, since we measure relative distance changes between the N- and C-termini of DP, our measurement of force-induced DP conformational changes remains valid irrespective of the tilted orientation of DP.

Furthermore, based on super-resolution microscopy, a model has been recently proposed where DP increases its tilt as desmosomes mature and become more adhesive, such that narrower and longer desmosomes correspond to stronger adhesive states [19, 38]. Since we observe narrower desmosomes in the K19-KO cells and wider desmosomes in WT cells, we assessed the intercellular adhesion strength of these cell lines using a dispase assay. In our assay, confluent sheets of either WT or K19-KO cells were detached using dispase II, a protease that gently cleaves cells from the substrate. Since cell-cell adhesion is mediated via both Ca^2+^-dependent classical cadherins (like E-cadherin) [39] and desmosomes which adhere in both a Ca^2+^-independent state and a Ca^2+^ dependent mode [40], we incubated the lifted cell sheets with 4 mM EGTA, a chelating agent for Ca^2+^, for 1 hour. Consequently, our assay selectively evaluated intercellular adhesion mediated mainly through the Ca^2+^-independent desmosomes. Following the EGTA treatment, the cell sheets were subjected to mechanical stress, resulting in the fragmentation of the sheets, as shown in Fig. S14A. Our results showed that the K19-KO cell sheets exhibited a significantly greater number of fragments than WT cell sheets. In contrast, K19-GFP cell sheets generated a similar number of fragments compared to WT cell sheets (Fig. S14B). These results suggest that in MCF7 cells, wider desmosomes do not correspond to weaker adhesive states. Furthermore, since the proposed model also suggests that desmosomes lengthen as they become more adhesive, we measured the DP length along the cell borders in in WT and K19-KO cells 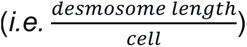. In contrast to the proposed model, our results showed a slight decrease in DP length in the weakly adhesive K19-KO cells compared to the strongly adhesive WT cells (Fig. S15). These results support our findings that changes in desmosome width are a direct result of DP conformational changes which are driven by actomyosin contractility and KIF organization. Our findings are also supported by a vast body of literature demonstrating that keratin deficiency is associated with impaired intercellular adhesion and increased cellular invasiveness [6, 41, 42].

Previous SMD simulations carried out on the long arm of the plakin domain (i.e., the crystal structure with PDB code 3R6N [24]) show that the SRs individually unfold when subjected to pulling force, which exposes the SH3 domain [43, 44]. Since our simulations are carried out on the entire plakin domain, which includes the flexible hinge between the long and short arms, we capture the transition of the plakin domain from a closed to an open conformation. The conformational change we measure is supported by previous SAXS measurements [17]. This transition of the plakin domain from a closed to an open structure also partially exposes the cryptic SH3 domain, which is known to act as a signal transduction adapter crucial in cell invasiveness, migration, and actin reorganization [45].

Many disease-causing mutations in DP are found within the plakin domain of the N-terminal region, affecting residues both hidden within the protein and exposed to the surrounding environment [46]. Specifically, mutations within the SR7-8 region of the plakin domain have been linked to arrhythmogenic ventricular cardiomyopathy (AVC) [16, 46]. Importantly, our simulations predict that the SR7-8 region participates in several key hydrogen bonds with SR5-6, which in turn stabilizes DP in its closed conformation. This raises the intriguing possibility that AVC mutations may interfere with the force-induced structural transition of DP from the closed to the open conformation.

## Materials and Methods

### Cell culture and stabilization

MCF7 WT cells (ATCC), MCF7 K19-KO cells, and MCF7 K19-GFP rescued cells were maintained in low-glucose Dulbecco’s Modified Eagle’s Medium (DMEM) (Gibco) supplemented with 10% Fetal Bovine Serum (FBS) (Life Technologies) and 1% penicillin-streptomycin (Sigma-Aldrich) in 5% CO2 at 37°C. The rescued cells were additionally treated with 100 ug/ml hygromycin for selection of K19-GFP. MDCK cells (ATCC) were cultured in low glucose DMEM (Gibco) cell culture media with 10% FBS and 1% PSK.

The K19-KO cells were generated using CRISPR/Cas9 as previously described [47]. We developed the K19-GFP rescue cell line following protocols described previously [48]. Briefly, K19-GFP was cloned into *K*19-KO cells from pEGFP-K19 plasmid into pLenti CMV Hygro (plasmid #17484. Addgene) using the following primers: forward 5’TCAGTCGACTGGATCCATGGTGAGCAAGGGCGAGG3’ and reverse 5’GAAAGCTGGGTCTAGTCAGAGGACCTTGGAGGC3’. Lentiviral supernatants were generated using pLenti-K19-GFP plasmid. Lentiviral supernatants collected 24h post-transfection were used to infect subconfluent K19-KO cells in three sequential 4 h incubations in the presence of 4 ug/ml polybrene (Sigma-Aldrich). Transductants were selected using a culture medium containing 100 ug/ml hygromycin beginning 48h after infection. The K19-GFP rescue cells were selected for further stabilization using fluorescence-activated cell sorting.

### Preparation of STED and confocal microscopy imaging samples

Approximately 500,000 cells were seeded for all the MCF7 cell lines, and approximately 250,000 cells were seeded for MDCK cells on collagen-coated coverslips (22 mm, No.1.5) in a single well of a six-well plate and were kept in 5% CO2 at 37°C for 24 h. A ‘calcium switch’ was then performed using the protocol described below to eliminate cell clusters and maintain a similar level of desmosome maturation across all samples. To perform a calcium switch, the cells were first rinsed with phosphate-buffered saline (PBS) (Gibco) solution at room temperature, followed by incubation in 4 mM EGTA (Millipore Sigma) and 10% low Ca^2+^ Fetal Bovine Serum (FBS) in Ca^2+^-free DMEM for 30 min. Following EGTA treatment, the cells were switched to a DMEM medium with 2 mM CaCl_2_ for 24 h to trigger the formation of mature desmosomes.

Next, cells were fixed either with ice-cold 1:1 methanol and acetone for 10 min (for keratin and DP staining) or with 3% PFA and 0.3% Triton X-100 for 10 min at room temperature (RT) (for F-actin staining). Samples were rinsed with PBS three times and blocked in 1% Bovine Serum Albumin (BSA) overnight at 4°C. Samples were then incubated with primary antibody (1:200) for 1 h at RT, followed by incubation with secondary antibody (1:600) for 30 min at RT. The samples stained with fluorescently labeled phalloidin (1:400) were incubated with phalloidin for 30 min at RT in the dark. Samples were washed three times with PBS after each antibody incubation. Lastly, samples were mounted to the glass slides using the ProLong Diamond Antifade Mountant (Life Technologies) and stored at 4°C in the dark.

## Drug treatments

Cells were cultured as previously described. For the blebbistatin treatment, cells were treated with 50 μM (±) blebbistatin (Millipore Sigma) for 24 h. Blebbistatin was dissolved in DMSO and diluted in the low-glucose DMEM (Gibco) supplemented with 10% FBS (Life Technologies) and 1% penicillin-streptomycin (Sigma-Aldrich) media. The control group was prepared with the same protocol but treated with 0.1% DMSO instead. For the Calyculin A treatment, cells were treated with 100 nM Calyculin A (Millipore Sigma) for 15 min. Calyculin A was dissolved in DMSO and diluted in the low-glucose DMEM (Gibco) supplemented with 10% FBS (Life Technologies) and 1% penicillin-streptomycin (Sigma-Aldrich) media. The control group was prepared with the same protocol but treated with 0.1% DMSO instead.

### Antibodies

The following primary antibodies were used: human anti-DSG2 (MAB947, R&D systems), rabbit anti-DPC antibody (A303-356A, Bethyl Lab), rabbit anti-DPN antibody (25318-1-AP, Proteintech), chicken anti-GFP (600-901-215, Rockland), mouse anti-K19 antibody (A53-B/A2) (Santa Cruz Biotechnology), mouse anti-K8 antibody (MA1-06318, ThermoFisher), mouse anti-K18 antibody (MA1-19047), control mouse IgG (sc-2025) (Santa Cruz Biotechnology), and mouse anti-actin antibody (66009-1-Ig, Proteintech). The following secondary antibodies are used: anti-chicken AF-488 (Invitrogen), anti-mouse AF-594 (Invitrogen), and anti-rabbit AF-647 (Invitrogen). Phalloidin AF-568 (Invitrogen) was used for F-actin staining.

The following antibodies were used for the Co-IP and western blot experiments: Anti-Keratin 19 antibody (A53-B/A2), anti-plakophilin-3 (E-10), anti-plakophilin-1 (10B2), anti-desmocolin-1 (A-4), anti-desmocolin-2/3 (7G6), anti-myosin-10 (A-3), and control mouse IgG (sc-2025) from Santa Cruz Biotechnology (Santa Cruz, CA), anti-Keratin 8 (17514-1-AP), ani-Keratin 18 (10830-1-AP), anti-actin antibody (66009-1-Ig), anti-desmoglein-2 (21880-1-AP), anti-plectin (29170-1-AP), anti-tubulin (66031-1-Ig), and anti-desmoplakin antibody (25318-1-AP) from Proteintech (Rosemont, IL).The following secondary HRP conjugated antibodies were used: anti-mouse-HRP (A4416), and anti-rabbit-HRP (A6154) from Sigma Aldrich (Louis, MO).

### STED imaging acquisition and processing

Immunofluorescence images were acquired at RT using a Leica TCS SP8 STED 3X confocal microscope (Leica Microsystems) equipped with a 100-x oil immersion objective (HC PL APO CS2), white light laser for excitation lines between 470-670 nm, and depletion laser lines at 592, 660, and 775 nm. The acquisition was performed using LasX software, followed by deconvolution in Huygens Pro software (Scientific Volume Imaging). Post-acquisition processing, including cropping and resizing, color change in the lookup table, adjustment of brightness and contrast, and inversion of grayscale images, was performed using ImageJ software.

### Confocal image acquisition and processing

Immunofluorescence images were acquired using a Leica Stellaris 5 confocal microscope (Leica Microsystems) equipped with a 63x oil immersion objective (HC PL APO CS2). Image acquisition was performed using LasX software, followed by post-processing using ImageJ. During post-processing, images were adjusted with inversion of grayscale, enhanced contrast with 0.3% saturated pixels, and reduced background noise using the despeckle functionality. All images were acquired using the same procedure.

### Distance measurement analysis

We used custom-made ImageJ macros and Python scripts to quantify the desmosome width along the entire cell-cell border. Since Dsg2 signals colocalize between DP “railroad” tracks, they were used as the reference to indicate desmosome location along the membrane. Briefly, Dsg2 signals were selected as regions of interest (ROIs), and an algorithm was used to determine the midline of the ROIs along the membrane axis and draw a segment along the midline every 200 nm to obtain the intensity profile of both Dsg2 and DPC or Dsg2 and DPN. Then, Bar, an ImageJ Plugin, was used to determine the location of peaks of different signals in their respective channels along the drawn segment. After obtaining the raw data, we used custom-made MATLAB code to calculate distances for full desmosome width represented by DPC-DPC or half desmosome width represented by Dsg2-DPC or Dsg2-DPN. To avoid false signals resulting from background noise or lines not fully drawn over the DP “railroad” track, we filtered out the data that showed less or more than two peaks in the DPC signals along the drawn segment or did not have the Dsg2 signal between the two DPC peaks. Furthermore, in certain desmosomes, the Dsg2 signals were observed as two peaks, as shown in Fig. S16A-B, implying that the desmosome had ruptured at the Dsg2 extracellular region. We eliminated these desmosomes from further analysis and only focused on intact desmosomes with unresolvable Dsg2 signals. For the DP line density measurement, we also used a custom-made ImageJ macro script for the quantitative analysis. Briefly, we first subtracted the background noise while retaining the DP signals along the cell borders. Then, we skeletonized these signals into segments to obtain the total amount of DP length along cell borders. Lastly, we obtained the DP line density by normalizing the total DP length using the number of cells in the image 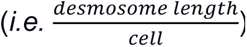.

### Co-immunoprecipitation

Co-immunoprecipitation was performed as described previously [49]. Cells grown on tissue culture plates were washed three times with 1X cold PBS. Cells were lysed at 4°C using ice-cold 1% triton lysis buffer (1% Triton X-100, 40mM HEPES (pH 7.5), 120mM NaCl, 1mM EDTA, 1mM phenylmethylsulfonyl fluoride, 10mM sodium pyrophosphate, 1 μg/ml each of chymostatin, leupeptin and pepstatin, 10 μg/ml each of aprotinin and benzamidine, 2 μg/ml antipain, 1mM sodium orthovanadate, 50mM NaF). Cell lysates were centrifuged to remove cellular debris and protein concentration was determined using Pierce BCA Protein Assay Kit and BSA as standards. Cell lysates were pre-cleared using Protein G Plus-Agarose beads (Santa Cruz Biotechnology) for 40 minutes at 4°C while rotating. While rotating, supernatants were incubated with the indicated antibody or IgG control for two hours at 4°C. Immuno-complexes were captured using Protein G Plus-Agarose beads and washed 4X using 1% triton lysis buffer supplemented with 300 mM NaCl.

### Gel electrophoresis and immunoblotting

Cells grown on tissue culture plates were washed three times with 1X cold PBS. Cells were lysed at 4°C using ice-cold 1% triton lysis buffer (1% Triton X-100, 40mM HEPES (pH 7.5), 120mM NaCl, 1mM EDTA, 1mM phenylmethylsulfonyl fluoride, 10mM sodium pyrophosphate, 1 μg/ml each of chymostatin, leupeptin and pepstatin, 10 μg/ml each of aprotinin and benzamidine, 2 μg/ml antipain, 1mM sodium orthovanadate, 50mM NaF). For immunoblotting, cell lysates were centrifuged at 14,000rpm for 5 minutes at 4°C to remove cellular debris. Protein concentration was determined using the Pierce BCA Protein Assay Kit following the manufacturer’s instructions (WG327385. Thermo Scientific). BSA (Equitech-Bio) prepared in 1% lysis buffer was used as the standard. Aliquots of protein lysates were resolved using SDS-PAGE and then transferred to nitrocellulose membranes (0.45um) (Amersham Protran; Cytiva) and immunoblotted using the indicated antibodies, followed by horseradish peroxidase-conjugated goat anti-mouse or goat anti-rabbit IgG (Sigma-Aldrich). Western Lightning Plus-ECL (Waltham, MA) or Amersham ECL Select Western was used as a blotting detection reagent. Signals were detected using iBright™ CL1500 Imaging System, Invitrogen (Thermo Fisher Scientific). The immunoblot result for comparing K19 levels among the three cell lines was conducted using a different method, as described previously [50].

### Structure preparation and MD simulation

Plakin domain structure (residues 178-950) was predicted using Alphfold2 [23]. Subsequently, the structure was prepared using PDBFixer [51] and missing hydrogens were added under pH 7.0 conditions. The protein N-terminus was designated with “NH3+” while “COO-” was assigned to the C-terminus. MD simulations were conducted on the FARM high-performance computing cluster at the University of California, Davis, with GROMACS 2022.3, as previously described [52–54]. The simulations were performed in the OPLS-AA/L force field [55] and TIP4P water model with a 10 Å radius cut-off for Van der Waals and electrostatic interactions. Electrostatic energy calculations employed the particle mesh Ewald method with a 0.16-grid spacing. At the beginning of the simulation, the plakin domain structure was positioned at the center of a dodecahedral box, ensuring a minimum distance of 1 nm from the boundary for every atom. The box was filled with water molecules and neutralized with charged ions (150 mM NaCl, 4 mM KCl, and 2 mM CaCl_2_). The system was relaxed with energy minimization and stabilized with equilibration under isothermal-isochoric and isothermal isobaric conditions using a modified Berendsen thermostat and Berendsen barostat. Post-stabilization, a 50 ns MD simulation was performed with 2-fs integration steps, maintaining a temperature of 300K using a v-rescale thermostat. Equilibration of the protein structure occurred within 20 ns. The number of hydrogen bonds observed between the long arm and the short arm was calculated using *gmx hbond*.

### Constant-force SMD simulations and analysis

The constant-force SMD simulations were performed as described previously [52–54]. The starting structures for the SMD simulations were the last frame of the corresponding MD simulations. The structures were placed at the side of a rectangular box (23 × 70 × 8 nm box, center at [11.5nm, 65nm, 4nm]). The system, containing ∼1,600,000 atoms, was relaxed and equilibrated under isothermal-isobaric conditions using the same protocol as in the MD simulation. The SMD simulations were performed at 310K temperature using a Nose-Hoover thermostat [56]. During each SMD simulation, we fixed the C-terminus of the plakin domain and pulled the N-terminal region (residues 919-950) along the longest axis of the box with a constant force ∼830 pN (500 kJ⋅mol−1⋅nm−1). The distances between the protein N-terminal and C-terminal were calculated using the *gmx pairdist*.

### Keratin filament orientation analysis

Confocal images were acquired as previously described for K8/DP and K18/DP staining. After image acquisition, we utilized a custom ImageJ macro script to determine the global dominant orientation of keratin filaments. Since filament orientation was defined as the angle between the keratin filament and the desmosome axis along the cell-cell junction, we first generated a mask based on DP signals outlining the cell border. This allowed us to define a region of interest (ROI) where the desmosome axis was set to 0°.

For each image, two rectangular ROIs were selected, each extending 2 µm from the cell border into the cytoplasm and having a length matching the desmosome mask. These ROIs were extracted, and the dominant orientation of keratin filaments was determined using the OrientationJ plugin. Specifically, OrientationJ computes the structure tensor, which captures intensity gradients at each pixel and extracts the principal direction of keratin filament orientation, represented as vector fields. The global dominant orientation was then derived by aggregating local orientations across the entire ROI. Consequently, each image provided two data points, each representing the global dominant orientation of desmoplakin-associated keratin filaments at the junction on either side of the cell-cell contact.

### Fluorescence lifetime imaging microscopy (FLIM)

Fluorescence lifetime data were acquired using a Leica TCS SP8 STED 3X confocal microscope, equipped with a pulsed white light laser (NKT Photonics) with wavelengths ranging from 470 to 670 nm, a 63x/1.35 NA oil immersion objective (HC PL APO CS2), and HyD detectors with photon counting capabilities. Images were captured at a scanning speed of 400 Hz with a resolution of 512 × 512 pixels. For each experimental condition, 10 images were acquired from 3 biological replicates.

Cells were cultured overnight on a 35 mm dish with a #1.5 glass-like polymer coverslip (Cellvis) to reach approximately 70% confluency. They were then transiently transfected with the DPI tension sensor, no-force control, and donor-only control constructs (Addgene), previously generated in an earlier study [28], using ViaFect Transfection Reagent (Promega) in reduced-serum Opti-MEM medium (ThermoFisher). After 8-hour incubation at 37°C with 5% CO₂, the culture medium was replaced with low-glucose DMEM (Gibco) supplemented with 10% FBS (Life Technologies) and 1% penicillin-streptomycin (Sigma-Aldrich). Cells were then treated with 1 µg/ml doxycycline to induce gene expression of the DPI constructs through a Tet-On system, which enables conditional gene expression in the presence of doxycycline by activating a doxycycline-sensitive transcriptional activator. The following day, cells were fixed with 4% PFA for 10 minutes at room temperature. After fixation, samples were maintained in Dulbecco’s PBS containing MgCl₂ and CaCl₂ (Sigma-Aldrich) and imaged at RT.

### FLIM-FRET analysis

Fluorescence lifetime data were analyzed using the built-in FLIM Wizard in LAS X software. For each lifetime image, 20–40 ROIs were manually selected, focusing on individual DP puncta. In each ROI, a bi-exponential decay model was fitted to the photon count time data to determine the fluorescence lifetime within each ROI. This model was chosen to mitigate the effects of the instrument response function (IRF) and autofluorescence, which typically result in small lifetime values. FRET efficiency (E) was then calculated using the donor lifetime in the presence of an acceptor (τ_DA_) and the mean donor-only lifetime (τ_D_), following the equation:

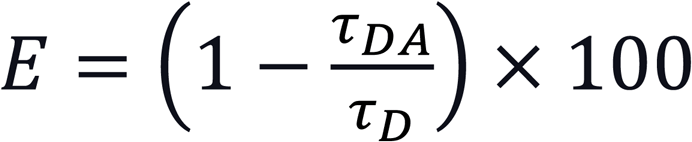

The mean donor-only lifetime was measured using donor-only constructs for each cell line, with WT cells exhibiting a mean lifetime of 2.771 ns and K19-KO cells showing a mean lifetime of 2.784 ns. To ensure reliable lifetime fitting, the minimum required photon count was set to 220 photons per image, as determined by the reduction in the spread of the fitted lifetime values.

### Statistical analysis

All statistical tests were performed using GraphPad Prism (GraphPad Software), and all graphs were plotted using custom-made Python code except those mentioned specifically. In the boxplots, the box represents the 25th and 75th percentiles with the median indicated, and whiskers reach 1.5 times the interquartile range (IQR), defined as the difference between the 25th and 75th percentiles. Data points outside the whiskers are shown as outliers. The number of independent experiments (N) and data points (n) is indicated in each figure legend. Data distribution was first tested based on Shapiro-Wilk’s test. Since most data was not normally distributed, non-parametric statistic tests were used for the analysis. Data were analyzed using Mann-Whitney’s U test. A multiple-group comparison was performed using the Kruskal-Wallis Test, followed by Dunn’s multiple comparison Test. Statistical tests used are specified in the figure legends. Statistical significance was represented at the level of *, P < 0.05; **, P < 0.01; ***, P<0.001; ns, P>0.05.

For the mean FRET efficiency analysis, statistical tests were conducted in R using a linear mixed-effects model, incorporating a grouped effect to account for puncta derived from the same image as described previously [28]. This approach corrects for statistical dependence among puncta within a single image. Specifically, the model assumes that grouped error across images follows a normal distribution centered around zero. By including this grouped effect, the analysis provides more conservative and robust statistical confidence compared to methods that treat each puncta as an independent data point. In R, the applied linear mixed-effects model was structured as *fretEfficiency ∼ isTensionSensor + (1|imageNumber)*, where *isTensionSensor* was assigned a value of 1 for tension sensor data and 0 for the corresponding truncated control data, while *imageNumber* uniquely identified each image.

## Acknowledgments

Research in the SS lab was supported by the National Institute of General Medical Sciences of the National Institutes of Health (R01GM121885). Research in the JSC lab was supported by the National Institute of General Medical Sciences of the National Institutes of Health (R15CA267890-01). STED imaging and FRET-FLIM imaging was performed at the UC Davis Advanced Imaging Facility. FACS was performed at the Flow Cytometry Shared Resource, funded by the UC Davis Comprehensive Cancer Center Support Grant (CCSG) awarded by the National Cancer Institute (NCI P30CA093373).

## Data availability

All data are made available in the manuscript and supporting information.

