## Supplemental Information for "Actomyosin forces trigger a conformational change in desmoplakin within desmosomes"

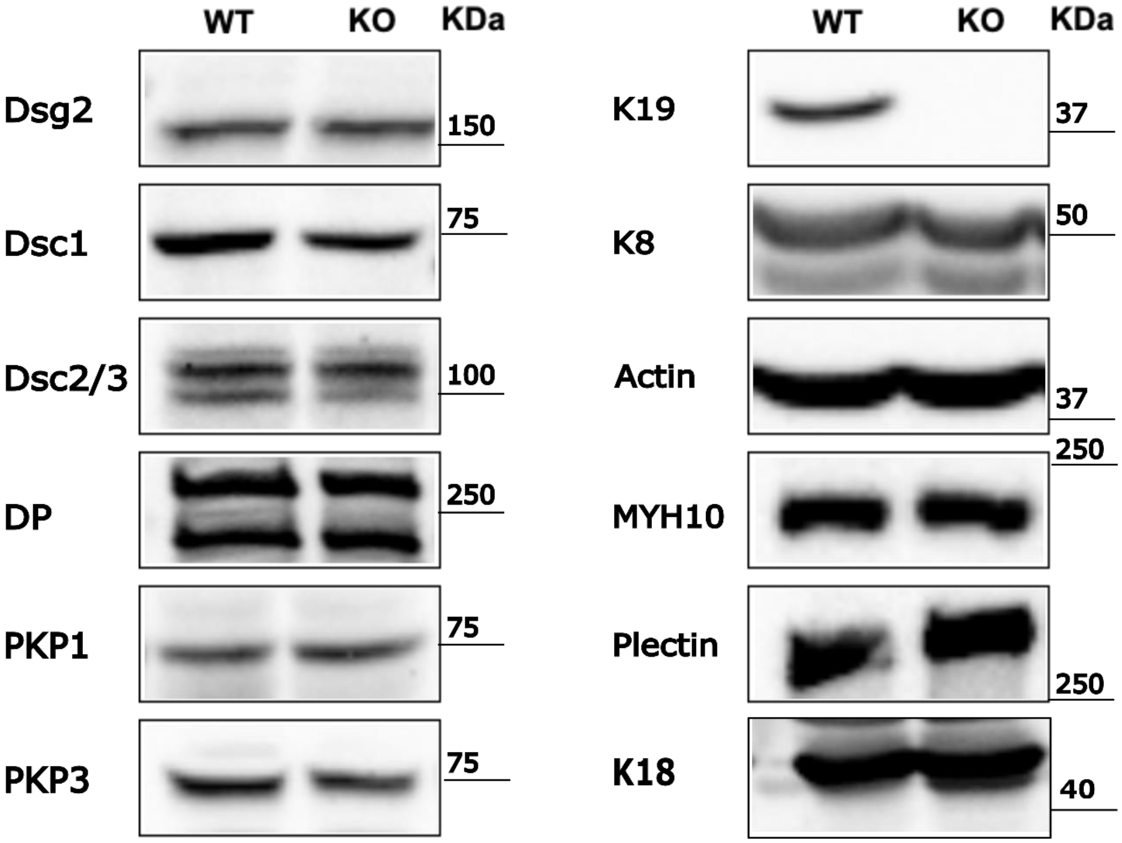


**Figure S1. Immunoblot of desmosomal cadherins, plaque proteins, and cytoskeletal proteins.** No discernable differences in cadherin, plaque protein, and actomyosin protein levels were detected between WT and K19-KO cells. K19 was present only in WT cells and not in K19-KO cells. Molecular weights from the corresponding protein ladders are marked. Dsg2: Desmoglien-2; Dsc1: Desmocolin-1, Dsc2/3: Desmocolin-2 and Desmocolin-3; DP: Desmoplakin; PKP1: Plakophilin-1; PKP3: Plakophilin-3; K19: Keratin-19; K8: Keratin-8; K18: Keratin-18. Immunoblotting was performed as described in the method section.

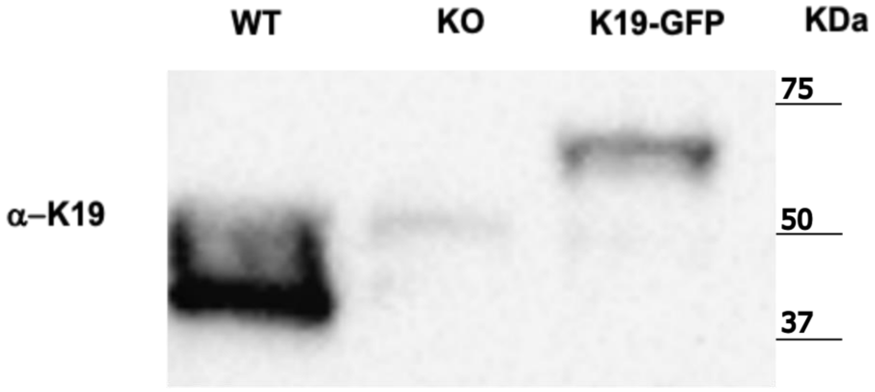


**Figure S2. Immunoblot results of K19 in WT, K19-KO, and K19-GFP cells confirms successful re-expression of K19 in the K19-GFP rescue cell line.** The K19 band in WT cells is ~ 3x the K19-GFP cells. Molecular weights measured from the ladder are indicated. K19 in WT cells has a molecular weight ~ 40 KDa while K19 tagged with GFP in K19-GFP cells has a molecular weight ~70 KDa. Immunoblotting was performed as previously described in the method and materials.

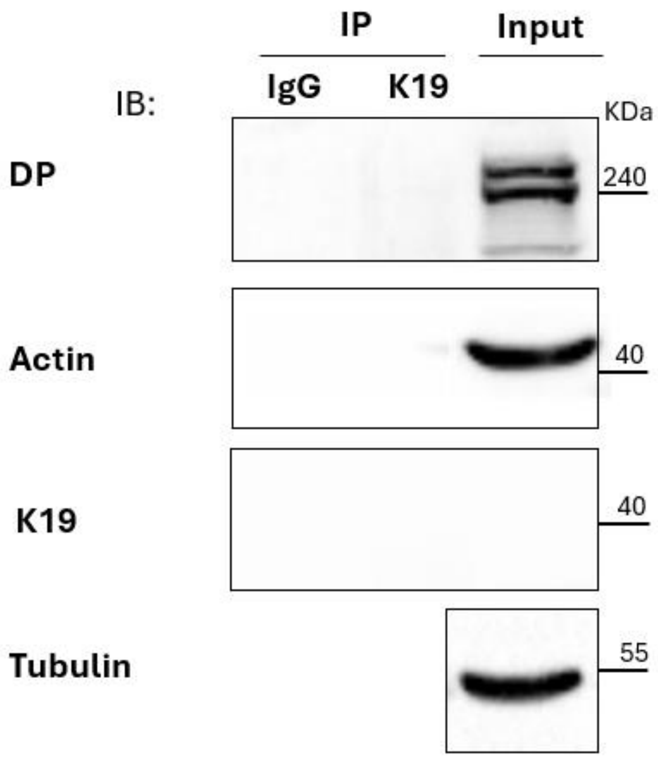


**Figure S3. Co-IP of K19 in K19-KO cells.** Co-IPs were performed with anti-K19 antibody or IgG control. The Co-Ips show that proteins pulled down with K19 in the WT were due to specific interactions.

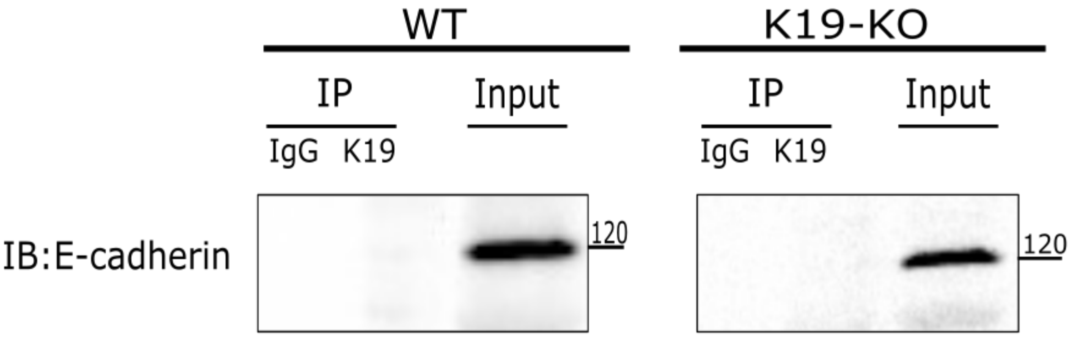


**Figure S4. Co-IP results for K19 show that the membrane-associated protein, E-cadherin, is not pulled down by K19.** No E-cadherin bands were observed for K19 in both WT and K19-KO cells, serving as an additional negative control to show that only specific interactions were observed for the Co-IP results for K19.

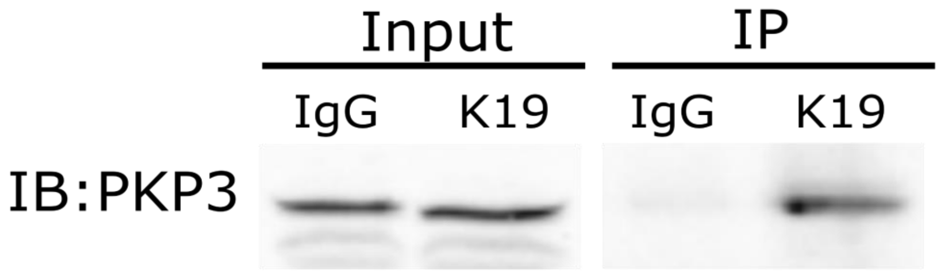


**Figure S5. Co-IP result for K19 shows an interaction between K19 and desmosomal plaque protein PKP3.** PKP3 bands were observed for both IgG control and K19 in the input samples, but the PKP3 band was only observed for K19 samples in the immunoprecipitation.

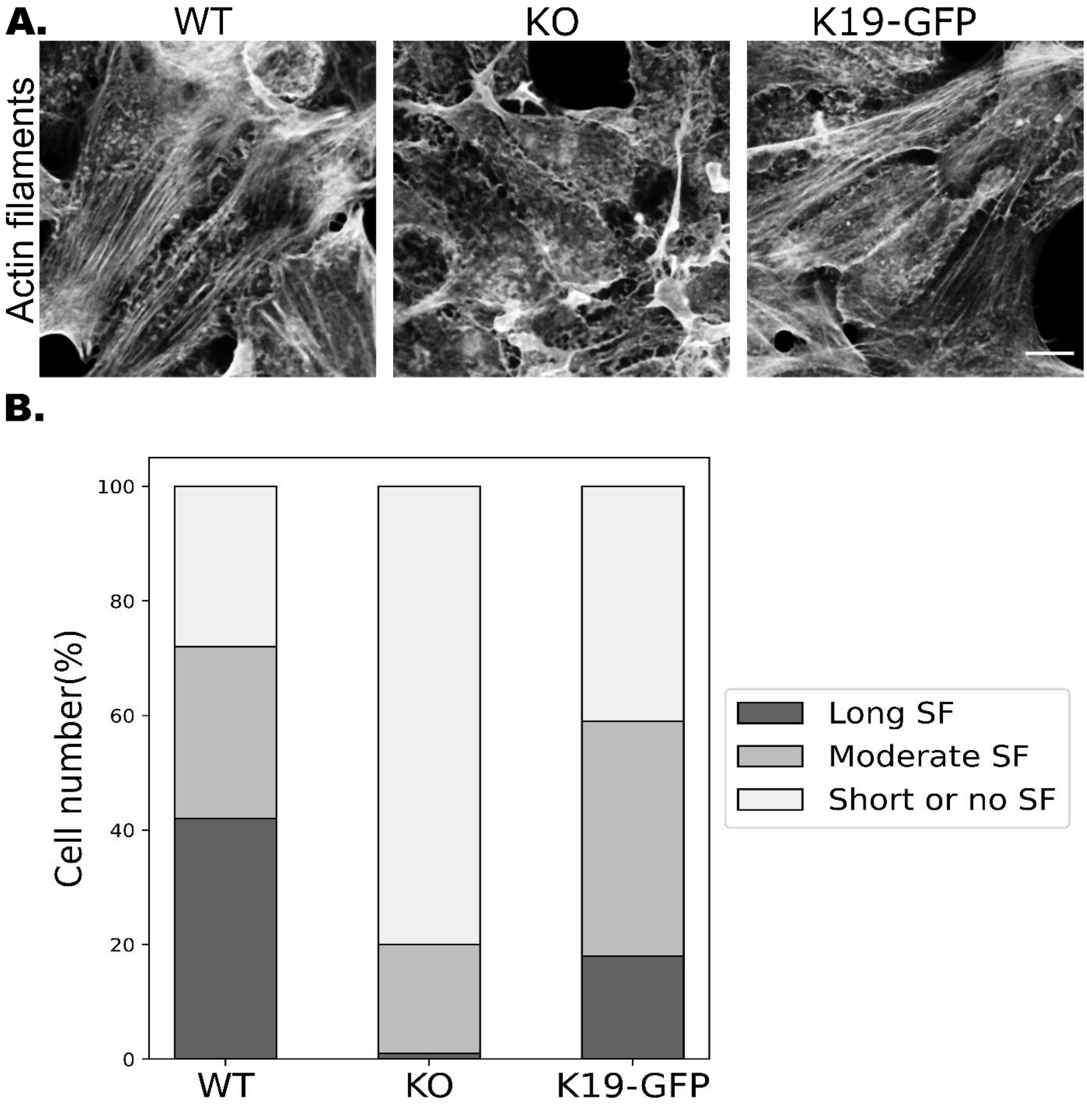


**Figure S6. Absence of K19 eliminates force-induced actin stress fiber formation. (A)** Confocal images of F-actin filaments in the WT, K19-KO, and K19-GFP rescued cell lines. Actin filaments were stained with Alexa568–phalloidin. Scale bar is 10 μm. **(B)** Quantitative analysis of actin stress fibers (SF) show a significant reduction of longitudinal actin stress fiber formation in the K19-KO cells compared to the WT and K19-GFP. Number of datapoints (n) = 461 (WT), 280 (K19-KO), 460 (K19-GFP); Number of replicates (N) = 3. The number of cells with long SF, moderate SF, and short or no SF was normalized by the total number of cells and shown as the percentage with respect to the total cell number in the plot.

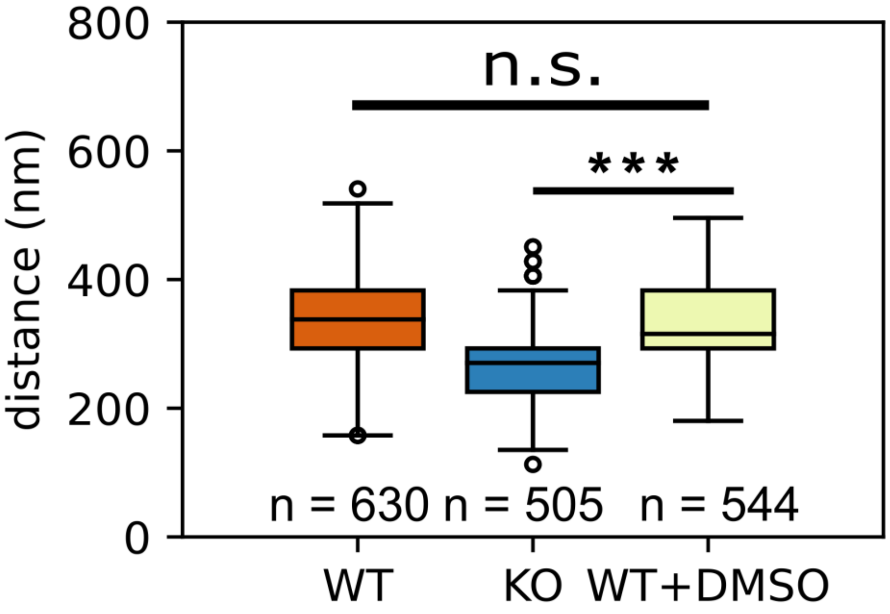


**Figure S7. Quantitative analysis of desmosome width shows no statistically significant differences between the DMSO control group and the WT cells.** Boxplots show the median, 25th, and 75th percentile with whiskers reaching the last data point; dots indicate the outliers in the data; n = 630 (WT), 505 (K19-KO), 544 (K19-GFP); N = 3. Kruskal-Wallis Test, followed by Dunn's multiple comparison Test; ***, P<0.001, ns, P>0.05.

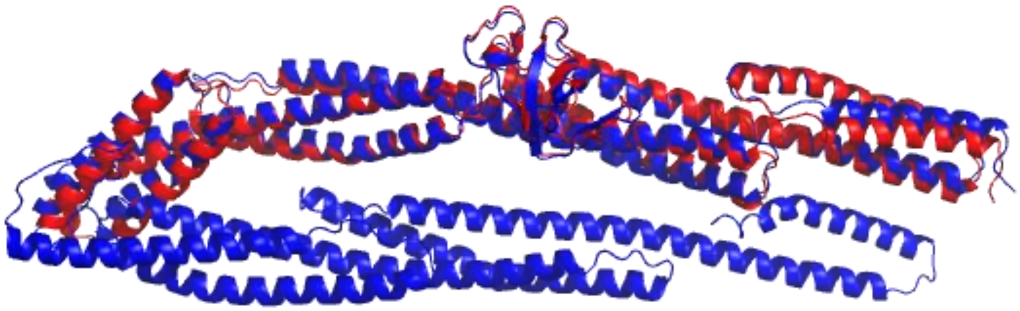

**Figure S8. Structural characterization of DP plakin domain.** Structural alignment between the crystal structure of the long arm of DP plakin domain (red, PDB code 3R6N) and the alpha-fold generated DP plakin domain suggests that the alpha-fold prediction is close to the real structure of DP plakin domain.

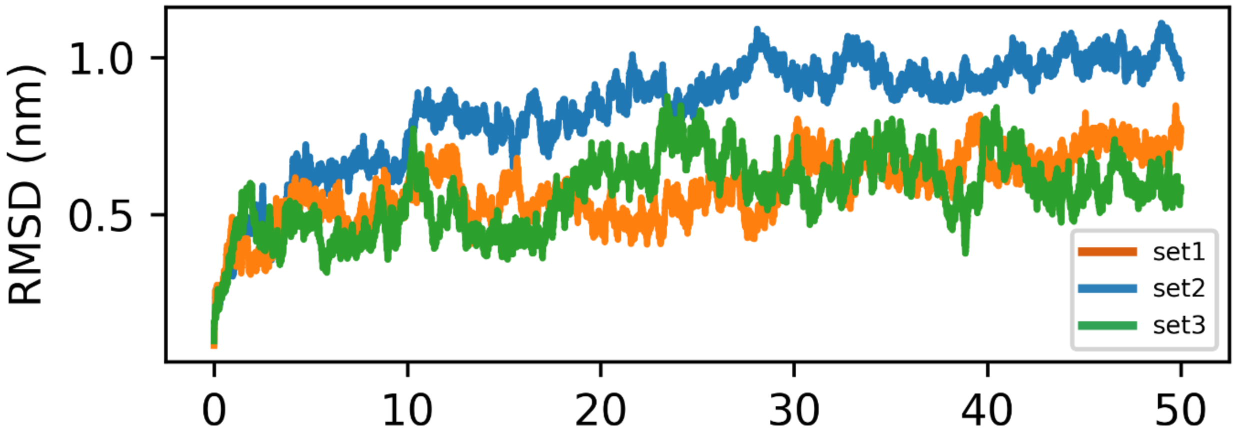


**Figure S9. Protein backbone RMSD in MD simulations relative to the initial structures at the start of simulation.** RMSD values were measured for three different sets. Stabilization of RMDS values within 10 ns suggest that structures are well equilibrated.

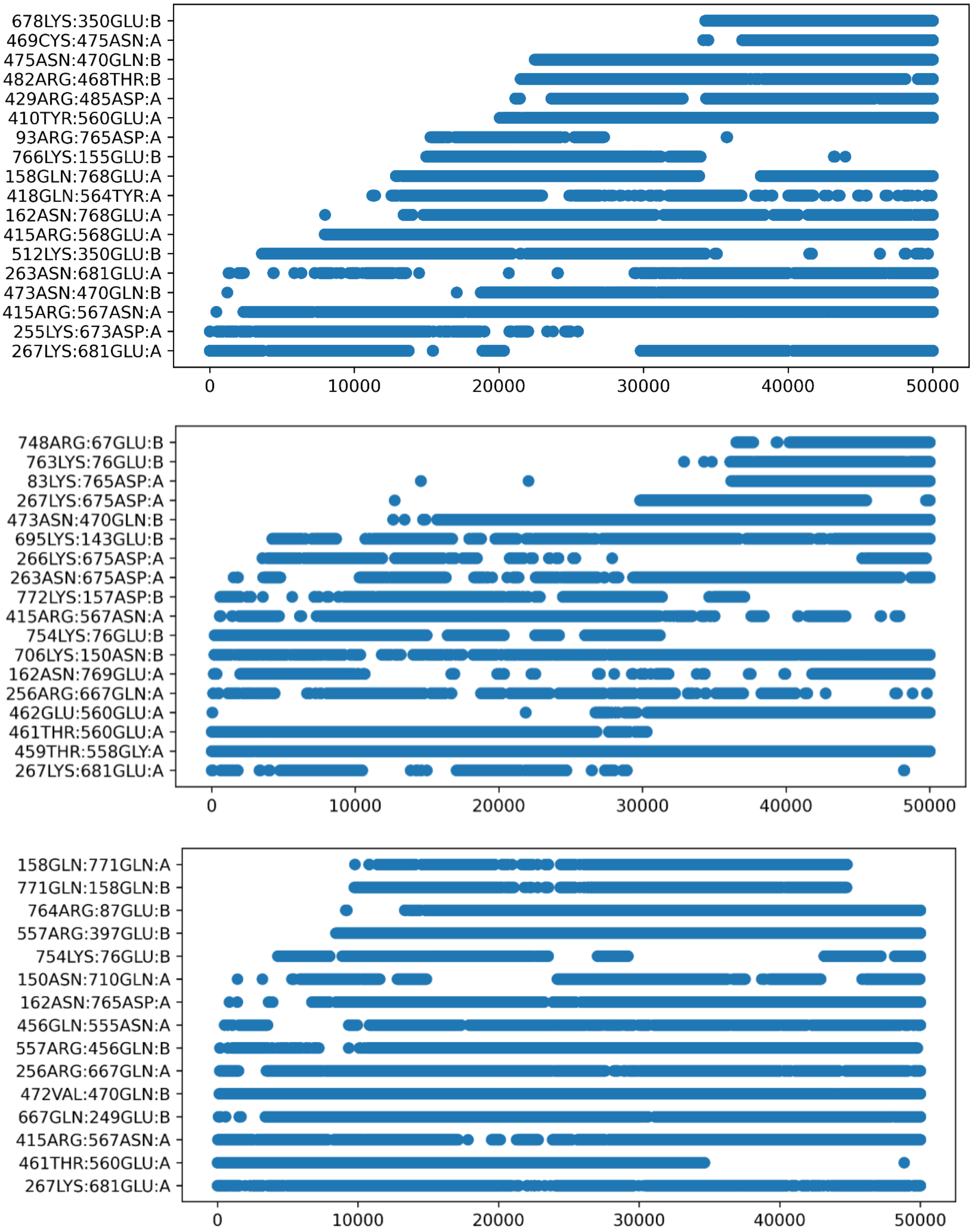


**Figure S10. Persistent hydrogen bonds (i.e. hydrogen bonds that last >25% of the time) are observed between the long arm (SR3-6) and the short arm (SR7-8 and SR8-CT) of the DP plakin domain.** Hydrogen bonds in set 1-3 are shown from top to bottom. Donor:acceptor pairs are labeled with "A", when hydrogen bond donor is from the long arm. Donor:acceptor pairs are labeled with "B", when hydrogen bond donor is from the short arm.

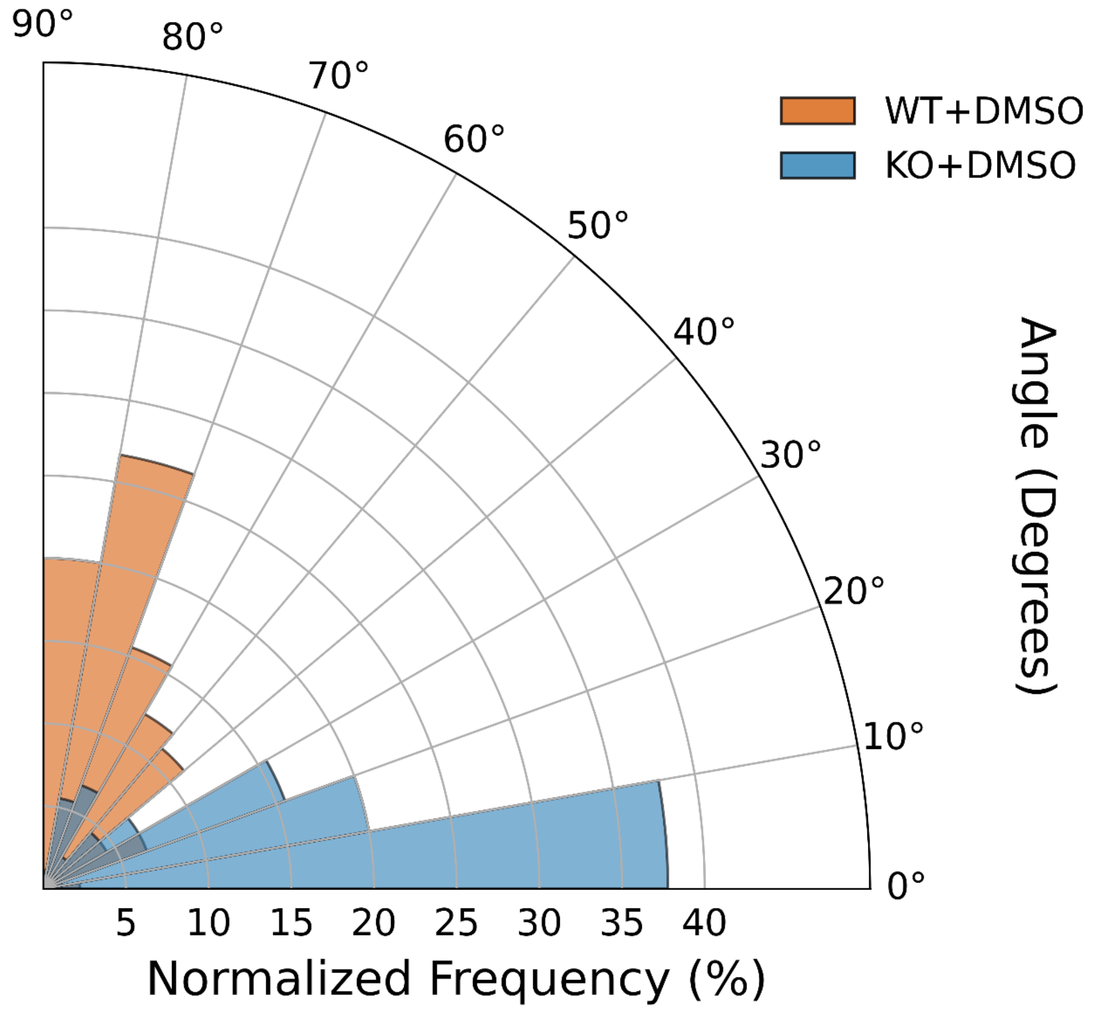


**Figure S11. Quantitative analysis of the global dominant orientation of K8 filaments in WT and K19-KO cells treated with DMSO.** Polar histogram of the global dominant orientation of K8 filaments in in WT and K19-KO cells treated with DMSO. n = 90 (WT+DMSO), 90 (KO+DMSO); N = 3. Mann-Whitney U test, $P < 0.0001$. The K8 filaments in WT+DMSO cells exhibit a more radial organization, while filaments in KO+DMSO cells are horizontally aligned. The results show similar keratin organization patterns between DMSO-treated and untreated cells (Fig. 4E), with WT and WT+DMSO cells both displaying primarily radial filament organization, and KO and KO+DMSO cells exhibiting predominantly horizontal filament alignment.

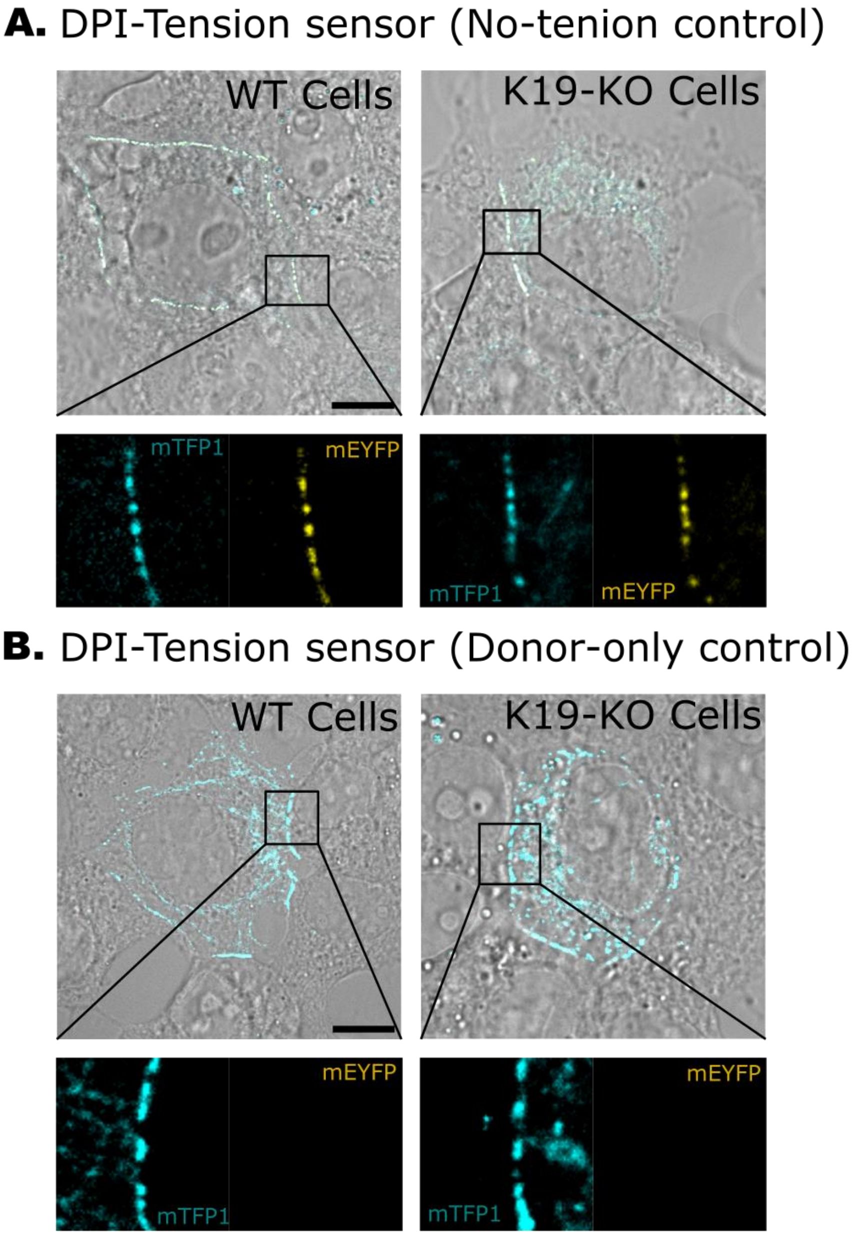


**Figure S12. Representative STED images of WT and K19-KO cells transfected with DPI-no-tension control (A) and the donor-only control (B)**. The overlay of fluorescent signals with the bright-field image shows the proper localization of DPI-no-tension control and DPI-donor-only control at the cell-cell contacts in both WT and K19-KO cells. Scale bar: 10 µm. The zoomed-in images reveal the formation of distinct DP puncta along the cell border.

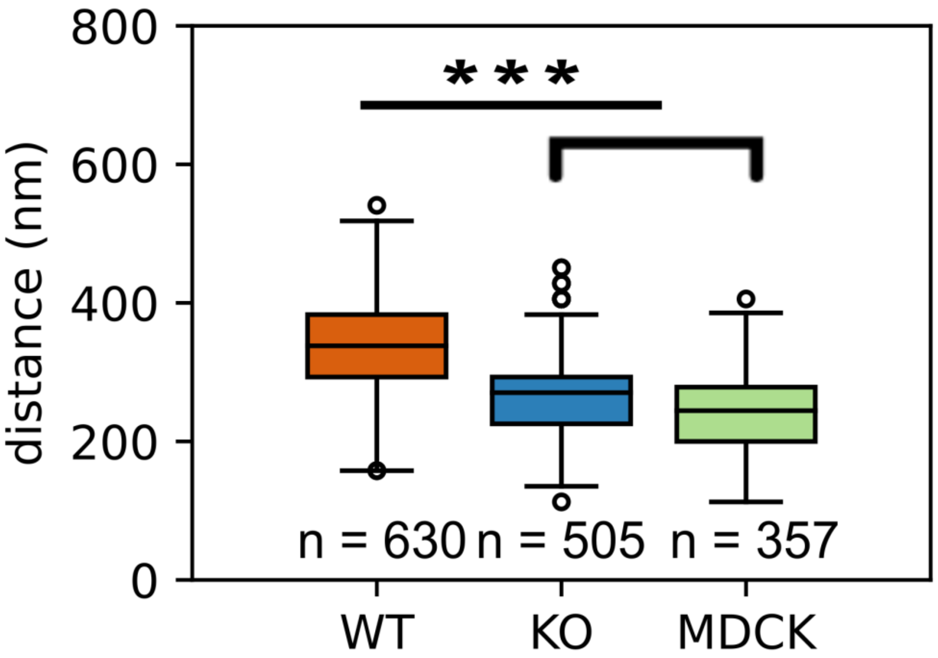


**Figure S13. Desmosome width in MDCK cells is lower than WT cells and comparable to K19-KO cells.** Boxplots show the median, 25th, and 75th percentile with whiskers reaching the last data point; dots indicate the outliers in the data; n = 630 (WT), 505 (K19-KO), 357 (MDCK); N = 3. Kruskal-Wallis Test, followed by Dunn's multiple comparison Test; ***, $P<0.001$.

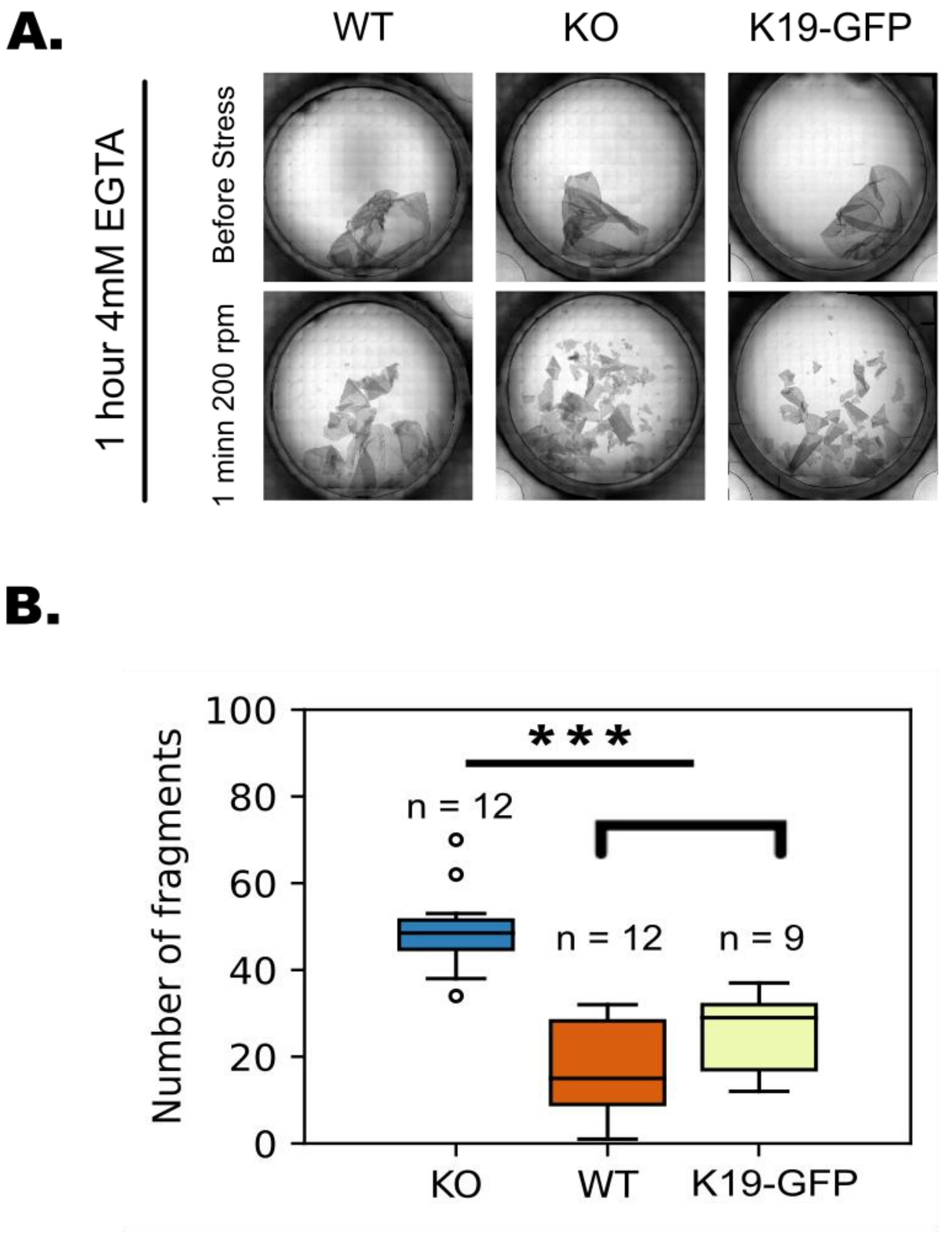


**Figure S14. Desmosome widening is not indicative of reduced adhesion strength. (A)** Dispase assay after 24 h plating. The confluent cell sheets are treated with 4 mM EGTA for 1 h. The images show the intact cell sheets for WT, K19-KO, and K19-GFP rescued cells before stress and fragmented cell sheets after applying mechanical stress. **(B)** Quantification of the dispase assay from K19-KO, WT, and K19-GFP cells. n = 12 (KO), 12 (WT), 9 (K19-GFP); N = 3. Kruskal-Wallis Test, followed by Dunn's multiple comparison Test; ***, P<0.001. The K19-KO cell sheets show a greater number of fragments compared to WT cell sheets, while K19-GFP cell sheets generate a similar number of fragments compared to WT.

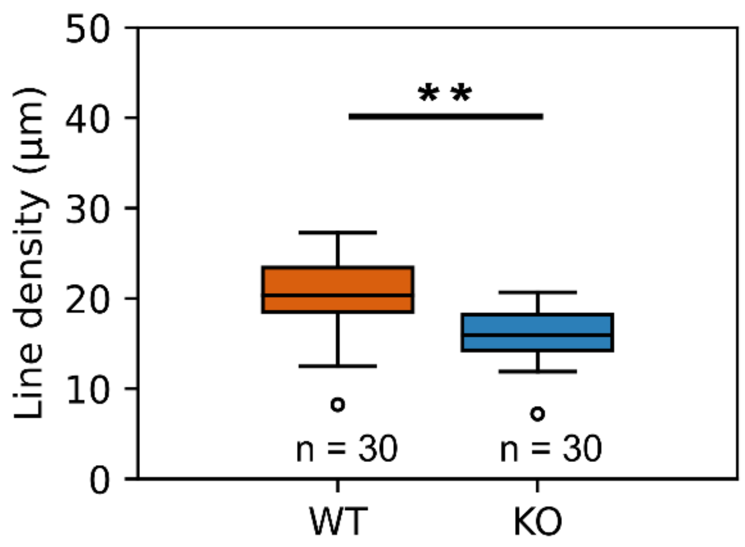


**Figure S15. The DP line density is marginally higher in the WT cells than the K19-KO cells.** Quantification of DP line density was shown to be marginally higher in the WT cells than in the K19-KO cells. The line density is defined as the desmosome length along the cell border normalized by number of cells. Number of datapoints (n) = 30 (WT), 30 (K19-KO); Number of replicates (N) = 3. Mann-Whitney's U test; **, P<0.01.

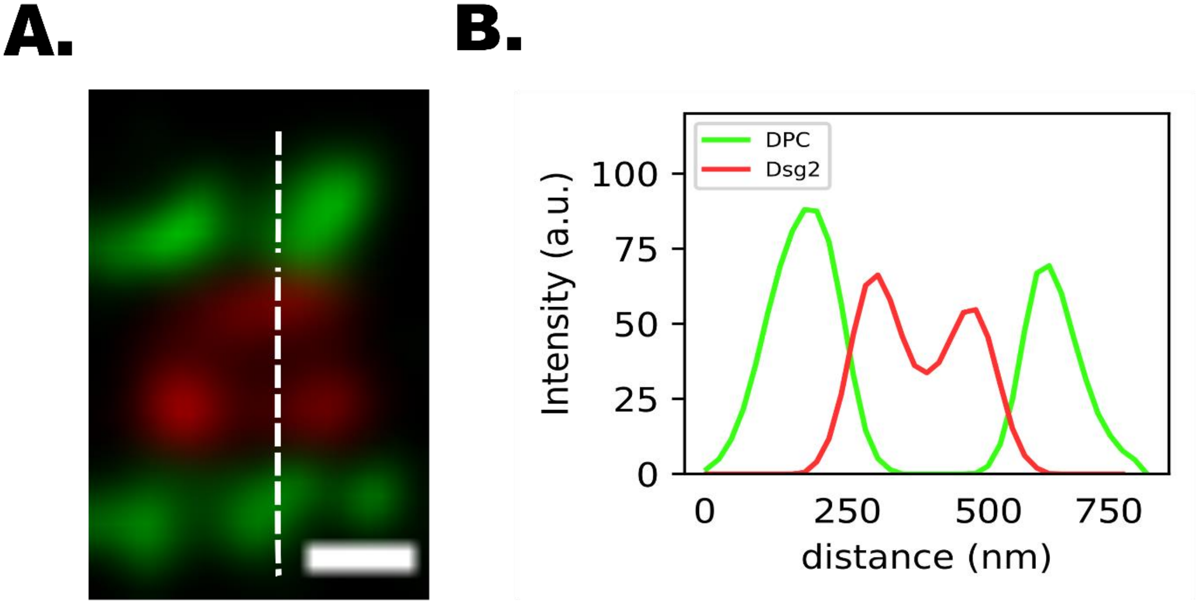


**Figure S16. Representative STED images of broken desmosomes in WT cells. (A)** DP railroad track immunolabeled for DPC (green) and Dsg2 (red). The Dsg2 signals are resolved (i.e. spatially separated) in the image. The scale bar is 200 nm. **(B)** Line-scan analysis of DPC and Dsg2 fluorescence intensity (indicated in A).

**Video S1. An example of constant-force SMD simulation, related to Fig. 3.** The constant-force SMD simulation illustrates the conversion of the DPN plakin domain from a folded (closed) to an extended (open) conformation upon pulling, which accounts for the elongation of 30-33 nm of the DP plakin domain. Color scheme: SR3-4 (green), SH3 (magenta), SR5-6 (blue), SR7-8 (orange), and SR8-CT (cyan).
